# Differential dynamics of microbial community networks help identify microorganisms interacting with residue-borne pathogens: the case of *Zymoseptoria tritici* in wheat

**DOI:** 10.1101/587667

**Authors:** Lydie Kerdraon, Matthieu Barret, Valérie Laval, Frédéric Suffert

## Abstract

**Background:** Wheat residues are a crucial determinant of the epidemiology of Septoria tritici blotch, as they support the sexual reproduction of the causal agent *Zymoseptoria tritici*. We aimed to characterize the effect of infection with this fungal pathogen on the microbial communities present on wheat residues, and to identify microorganisms interacting with it. We used metabarcoding to characterize the microbiome associated with wheat residues placed outdoors, with and without preliminary *Z. tritici* inoculation, comparing a first set of residues in contact with the soil and a second set without contact with the soil, on four sampling dates in two consecutive years.

**Results:** The diversity of the tested conditions, leading to the establishment of different microbial communities according to the origins of the constitutive taxa (plant only, or plant and soil), highlighted the effect of *Z. tritici* on the wheat residue microbiome. Several microorganisms were affected by *Z. tritici* infection, even after the disappearance of the pathogen. Linear discriminant analyses and ecological network analyses were combined to describe the communities affected by infection. The number of fungi and bacteria promoted or inhibited by inoculation with *Z. tritici* decreased over time, and was smaller for residues in contact with the soil. The interactions between the pathogen and other microorganisms appeared to be mostly indirect, despite the strong position of the pathogen as a keystone taxon in networks. Direct interactions with other members of the communities mostly involved fungi, including other wheat pathogens. Our results provide essential information about the alterations to the microbial community in wheat residues induced by the mere presence of a fungal pathogen, and vice versa. Species already described as beneficial or biocontrol agents were found to be affected by pathogen inoculation.

**Conclusions:** The strategy developed here can be viewed as a proof-of-concept focusing on crop residues as a particularly rich ecological compartment, with a high diversity of fungal and bacterial taxa originating from both the plant and soil compartments, and for *Z. tritici*-wheat as a model pathosystem. By revealing putative antagonistic interactions, this study paves the way for improving the biological control of residue-borne diseases.

## Background

Septoria tritici blotch (STB) is one of the most important disease of wheat (*Triticum aestivum*), causing yield losses averaging 20% on susceptible wheat varieties and 5–10% on wheat varieties selected for disease resistance and sprayed with fungicide in Northwestern Europe [1]. It is caused by the hemibiotrophic, heterothallic, ascomycete fungus *Zymoseptoria tritici* [2], which initiates its sexual reproduction on senescent tissues [3]. STB is clonally propagated between wheat plants during the cropping season by pycnidiospores (asexual spores), which are splash-dispersed upwards over short distances. Wind-dispersed ascospores (sexual spores), mostly produced on wheat residues, initiate subsequent epidemics. Thus, wheat residues are a crucial, but often neglected determinant of the epidemiology of STB during the interepidemic period, as they support the sexual reproduction of the pathogen, maintaining diversity within populations and influencing adaptive dynamics in response to selection pressures [4], through the rapid evolution of fungicide resistance [5–8] or the breakdown of wheat resistance genes [9], for example.

The identification of microorganisms interacting with pathogens is an increasingly important issue for both academic and operational research on the development of biological control solutions [10,11]. In plant, animal and human epidemiology, increasing numbers of studies are trying to characterize variant microbial populations associated with specific disease stages, or temporal changes in the microbial populations during disease progression [12–14]. The pathogen and its cohort of associated microorganisms, which may influence its persistence, transmission and evolution, are together known as the “pathobiome” [15]. Pathobiome research has advanced significantly with the advent of high-throughput sequencing technologies, which have made it possible to describe and follow the diversity of the microbial communities associated with the pathogen during its life cycle, during both the epidemic and interepidemic periods.

The dynamics of microbial communities have been studied in detail during the vegetative and reproductive stages of the plant life cycle, but very few studies during and after plant senescence (e.g. [16,17]). The specific, central position of crop residues in agrosystems was long neglected, but these residues should be seen as both a fully-fledged matrix and a transient compartment: a compartment originating from the plant (temporal link), then in close contact with the soil (spatial link), with variable rates of degradation over the following cropping season, according to the plant species, the cropping practices used, and the climatic conditions in the year concerned [16,19–21]. In addition, the rare studies focusing on the evolution of microbial communities in crop residues performed to date were conducted in microcosms, with sterilized residues (e.g. [22]), in which this compartment is much less complex than under natural conditions.

Several studies have investigated the potential beneficial effects of microorganisms for limiting the development of a plant pathogen during its saprophytic stage on natural crop residues (e.g. *Aureobasidium pullulans* and *Clonostachys rosea* inhibiting the sexual stage of *Didymella rabiei* on chickpea residues [23]; *Trichoderma harzianum* [24,25], *Microsphaerelopsis* sp. [26], *C. rosea* [27,28] and *Streptomyces* sp. [29] reducing *Fusarium graminearum* inoculum (perithecia, the sexual fruiting bodies) on wheat or maize residues, as summarized in [30]). Other studies have focused on the general impact of cropping practices, such as the increase in microbial soil antagonists induced by the addition of green manure to the soil (e.g. [19,31]). Some phyllosphere microorganisms selected for their antifungal activity against *Z. tritici* (*Bacillus megaterium* [32]; *Pseudomonas fluorescens* [33]; *Cryptococcus* sp., *Rhodotorula rubra* and *Penicillium lilacinum* [34]; *T. harzianum* [35]; Trichoderma koningii [36]) have been tested *in planta* against the asexual, pathogenic stage of the pathogen (typically on wheat seedlings), but not against the pathogen during its sexual, saprophytic stage. Moreover, no microbial antagonists of *Z. tritici* have been isolated from wheat residues, despite the dense population of this habitat with a high diversity of microbial taxa [16].

The taxonomic structure of microbial communities associated with maize [17] and wheat [16] residues has recently been described under natural conditions. In addition to *Z. tritici*, the microbial communities associated with wheat include *Clonostachys* sp., *Aureobasidium* sp., *Chaetomium* sp. and *Cryptococcus* sp. [16], all of which are potential competitors. However, the presence of microorganisms in the same ecological niche, as highlighted in such descriptive approaches, does not necessarily mean that interactions actually occur between them. Many other non-interacting microorganisms (pathogens, endophytes) are also present on the residues. Moreover, microbial communities change during the physical degradation of the residues, probably modifying interactions between microorganisms over time [16]. Ecological network analysis has made it possible to detect putative interactions between microorganisms. For instance, Jakuschkin *et al*. [13] detected significant changes in foliar fungal and bacterial communities following the infection of pedunculate oak with *Ersysiphe alphitoides* (the causal agent of oak powdery mildew), and Cobo-Diaz *et al*. [17] identified candidate antagonists of toxigenic *Fusarium* spp. among the species present in maize residues. The use of co-occurrence networks in these two studies highlighted a set of bacteria and fungi that might be useful for managing plant pathogens.

In this study, our goal was to identify fungi and bacteria potentially interacting with *Z. tritici* during its sexual reproduction on wheat residues. To this end, we compared the structure of microbial communities associated with wheat residues with and without *Z. tritici* inoculation, by metabarcoding, combining linear discriminant analyses (LDA) and ecological network analyses (ENA). The response of microbial communities to *Z. tritici* infection was assessed during the interepidemic period between two successive crops, for two sets of wheat residues, one left outdoors in contact with the soil, and the other left outside but not in contact with the soil, at different sampling dates during two consecutive years. The diversity of experimental conditions was expected to lead to the establishment of different microbial communities according to the origin of the constitutive taxa (plant or soil), thereby increasing the probability of detecting effects of *Z. tritici* on the residue microbiome, and of the residue microbiome on *Z. tritici*.

## Results

### Overall diversity of the bacterial and fungal communities on residues

The response of the residue microbiome to *Z. tritici* inoculation was assessed by analyzing the composition of the fungal and bacterial communities of wheat residues, after inoculation with *Z. tritici* (n=240) or in the absence of inoculation (n=240). We also investigated the impact of cropping season (n=2), season (n=4), and soil contact (n=2) on the dynamics of these communities (see materials and methods for a detailed explanation of the experimental design; Figure 1).

**Figure 1.**
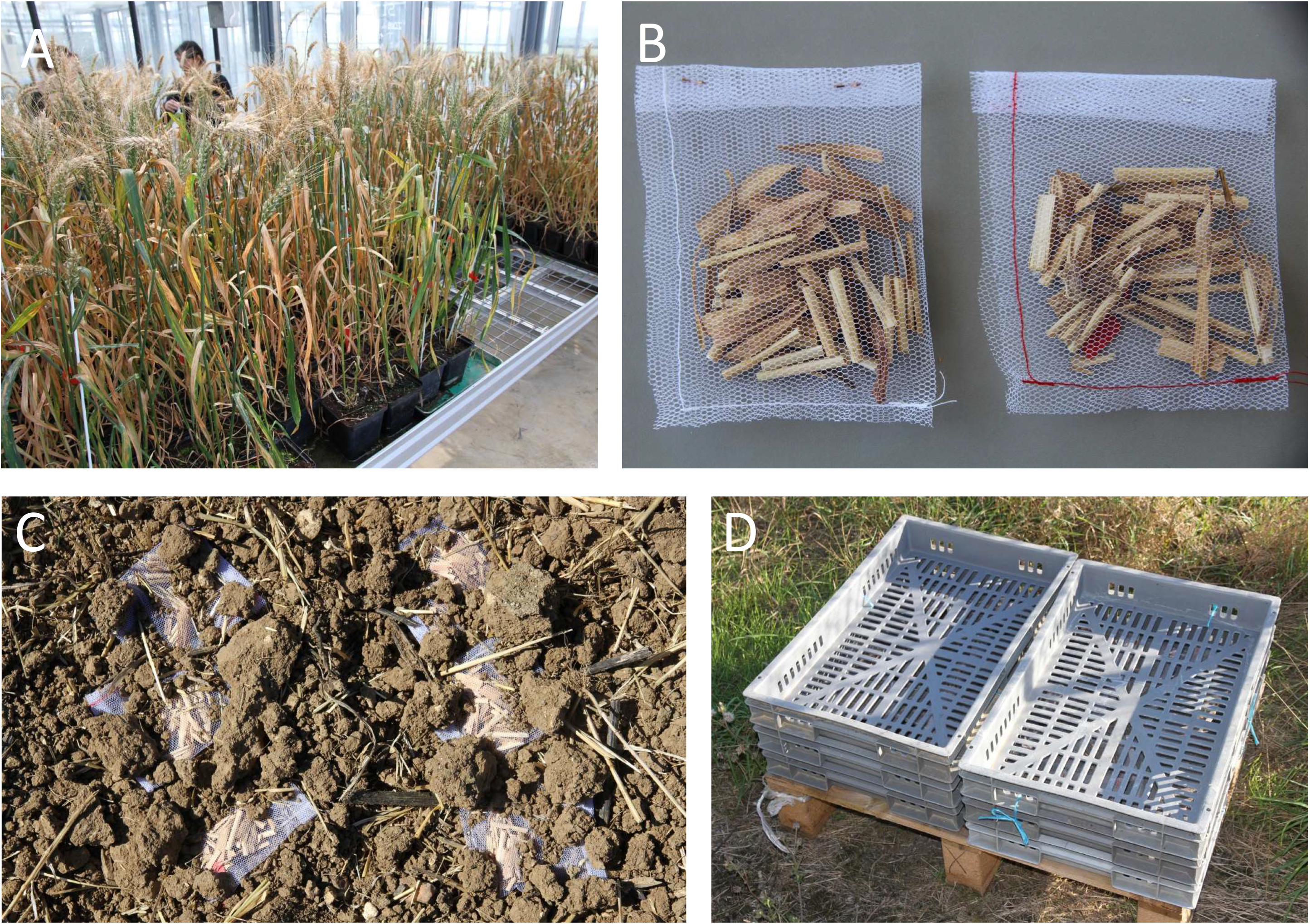
Preparation of wheat residues. (A) Adult wheat plants were inoculated with *Zymoseptoria tritici* under greenhouse conditions. (B) Sealed nylon bags containing wheat residues, consisting of stem and leaf fragments of approximately 2 cm in length (red yarn for residues from wheat plants inoculated with *Z. tritici*; white yarn for those from non-inoculated plants). (C) “Soil contact” treatment: nylon bags were left on the ground of the field and partially covered with soil (one of the 15 sampling points). (D) “Above ground” treatment: plastic grids containing nylon bags placed outside the field.

We investigated the structure of the residue microbiome by analyzing the v4 region of the 16S rRNA gene and ITS1. Overall, 996 bacterial amplicon sequence variants (ASVs) and 520 fungal ASVs were obtained from 390 and 420 samples, respectively. Some samples (July 2016) were removed from the analysis due to the co-amplification of chloroplasts.

The high relative abundance (RA) of ASVs affiliated to *Zymoseptoria* in samples collected in July 2016 (21.5±9.8%) and 2017 (30.3±7.1%) highlights successful colonization of the wheat tissues by this pathogen following inoculation (Figure 2). However, the RA of *Zymoseptoria* rapidly decreased to 2±1.64% and 1.4±0.9% on residues not in contact with the soil (above ground residues) collected in October 2016 and 2017, respectively, and this species was below the limit of detection in December and February. For residues in contact with soil, this decrease occurred more rapidly, with *Zymoseptoria* ASV already undetectable in samples collected in October.

**Figure 2.**
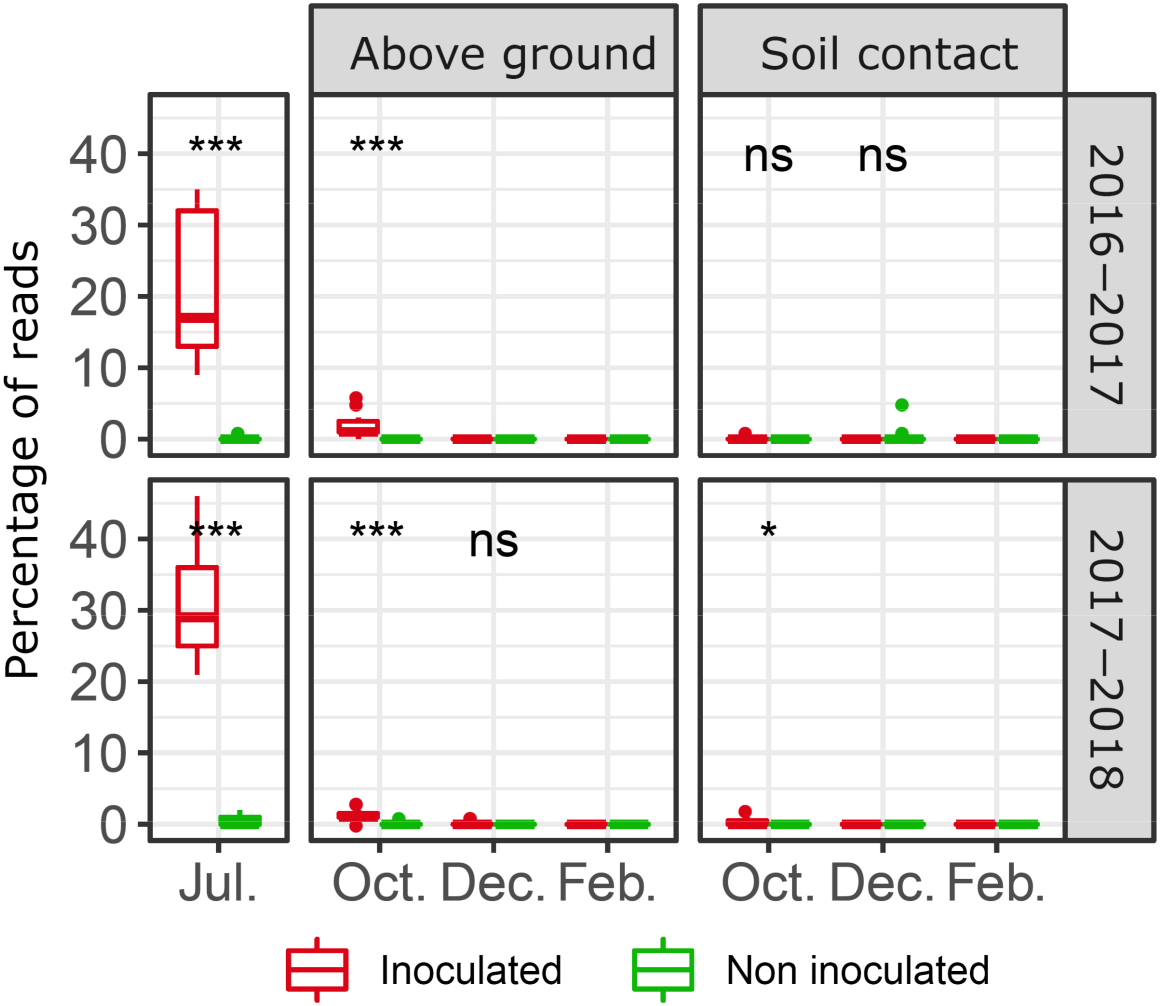
Relative abundance of *Zymoseptoria tritici*. Each box represents the distribution of the relative abundances of genera for the 15 sampling points. Wilcoxon tests were performed for inoculation condition (NS: not significant; * *p*-value<0.05; ** *p*-value <0.01; *** *p*-value <0.001).

Alpha diversity, estimated with the Shannon index, was low in July for both bacterial (2.70±0.75) and fungal communities (1.82±0.19; Suppl. Figure 1). A gradual increase was then observed during residue degradation. *Z. tritici* inoculation had no impact on bacterial alpha-diversity, but decreased fungal diversity (Kruskal-Wallis: *p* = 0.008). More specifically, bacterial diversity was higher in inoculated residue samples in July 2017 (2.92±0.80 for inoculated samples versus 2.47±0.6 for non-inoculated samples; Wilcoxon: *p* = 0.022), but no such difference was detected for the other sampling dates. Conversely, for fungal communities, inoculation had no effect in July, but led to a significant decrease in diversity in subsequent months during the second cropping season (October and December 2017, for the two soil contact conditions).

Beta diversity analysis (Bray-Curtis index) showed large dissimilarities between bacterial community composition in July and at the other sampling dates, as illustrated in the hierarchical clustering of the samples, justifying separate analyses and MDS representations (Figure 3). Inoculation with *Z. tritici* had a minor effect on bacterial communities (Table 1), with only 11.5% of the variance explained for samples collected in July (PERMANOVA: *p* = 0.001). By contrast, in the same month, inoculation was the structuring factor for fungal communities, accounting for 33.3% of the variance (PERMANOVA: *p* = 0.001). For subsequent samplings (October, December and February), temporal conditions (seasonality and cropping season) were the main factors influencing fungal communities. Soil contact was the main structuring factor for bacterial communities, with a stronger effect than seasonality or cropping season (Table 1).

**Figure 3.**
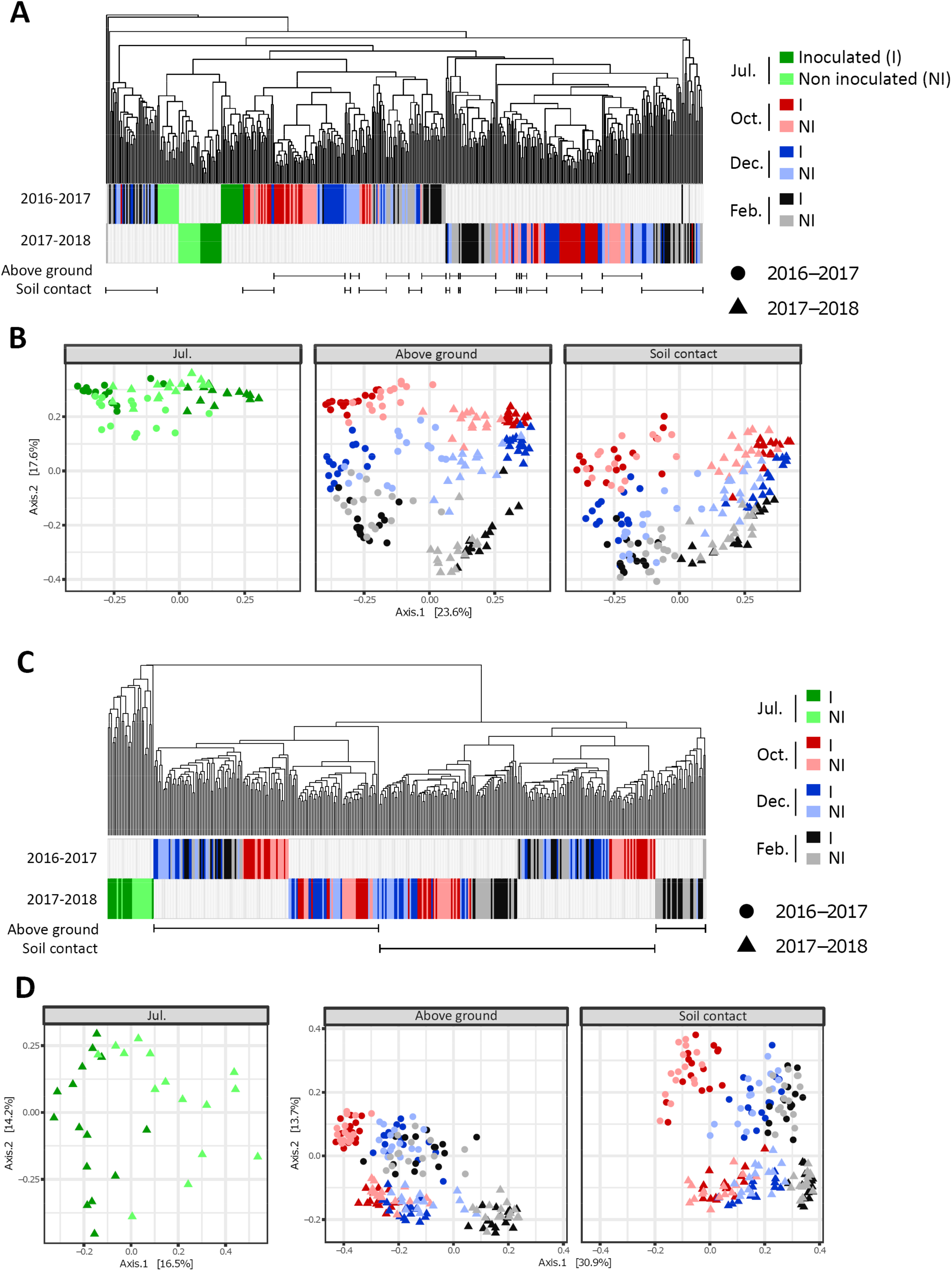
Dissimilarities between microbial communities. Beta diversity analyses for fungal (A, B) and bacterial (C, D) communities originating from 420 wheat residue samples. Hierarchical clustering (A, C) and multidimensional scaling (B, D) are based on the compositional distances between samples (Bray Curtis distance matrix). (A, C) Visualization of compositional distances between samples through hierarchical clustering with the average linkage method. The samples (15 sampling points per treatment) corresponding to the two cropping seasons (year) are represented by the two colored horizontal series (2016-2017, 2017-2018). Effects of seasonality are highlighted by different colours, corresponding to the different sampling dates (July: green; October: red; December: blue; February: gray). The intensity of the colors distinguishes between samples obtained from plants inoculated with *Z. tritici* (I, dark hues) and non-inoculated samples (NI, light hues). “Above ground” and “soil contact” treatments are represented by horizontal lines, with each sample considered separately. (B, D) Visualization of compositional distances between samples through multidimensional scaling (MDS). Each data point corresponds to one sample of wheat residues. The shape of the points (circles: 2016-2017; triangles: 2017-2018) corresponds to the cropping season (year effect); the colors, similar to those used in graphs A and C, correspond to the sampling dates (seasonality effect). For fungal communities, MDS analysis was performed on all samples together, whereas for bacterial communities, the analyses of the July samples and samples from all other sampling dates (October, December, and February) were separated, in accordance with the large differences between the communities of these samples shown in the clustering analysis (C). For a sake of clarity, the MDS are shown according to the soil contact condition.

**Table 1.** Results of PERMANOVAs analyzing the effects of cropping season, sampling date, contact with soil and inoculation on fungal and bacterial communities. Factors were tested with the adonis2 function of the vegan package. PERMANOVAs were performed with all tested factors together, with the “margin” option.

|  |  | Tested factors | Explicated variability | p-value |
| --- | --- | --- | --- | --- |
| Fungi | July | Season | 0.197 | 0.001 |
|  |  | Inoculation | 0.333 | 0.001 |
|  | Oct - Dec - Feb | Season | 0.217 | 0.001 |
|  |  | Sampling dates | 0.136 | 0.001 |
|  |  | Contact with soil | 0.096 | 0.001 |
|  |  | Inoculation | 0.012 | 0.001 |
| Bacteria | July | Season | – * | – * |
|  |  | Inoculation | 0.115 | 0.001 |
|  | Oct - Dec - Feb | Season | 0.128 | 0.001 |
|  |  | Sampling dates | 0.168 | 0.001 |
|  |  | Contact with soil | 0.195 | 0.001 |
|  |  | Inoculation | 0.006 | 0.001 |

\* Not tested. Due to the larger proportion of chloroplast sequences among the 16S rRNA gene products obtained from living plant tissues compared to dead tissues, all samples from July 2017 were removed from the analysis.

### Impact of contact with the soil on microbial communities

The significant impact of soil contact on microbial communities highlighted differences in the process of wheat residue colonization. MDS analysis suggested that the communities of “above ground” residue samples collected in October were less different from those collected in July than from the communities of “soil contact” samples also collected in October (Figure 4). Contact with the soil, therefore, caused a greater change in communities, suggesting competition between plant-associated taxa and soil-borne taxa. Taxonomic differences between the communities present on residues in contact with the soil and those present in above ground residues were highlighted in linear discriminant analysis (LDA).

**Figure 4.**
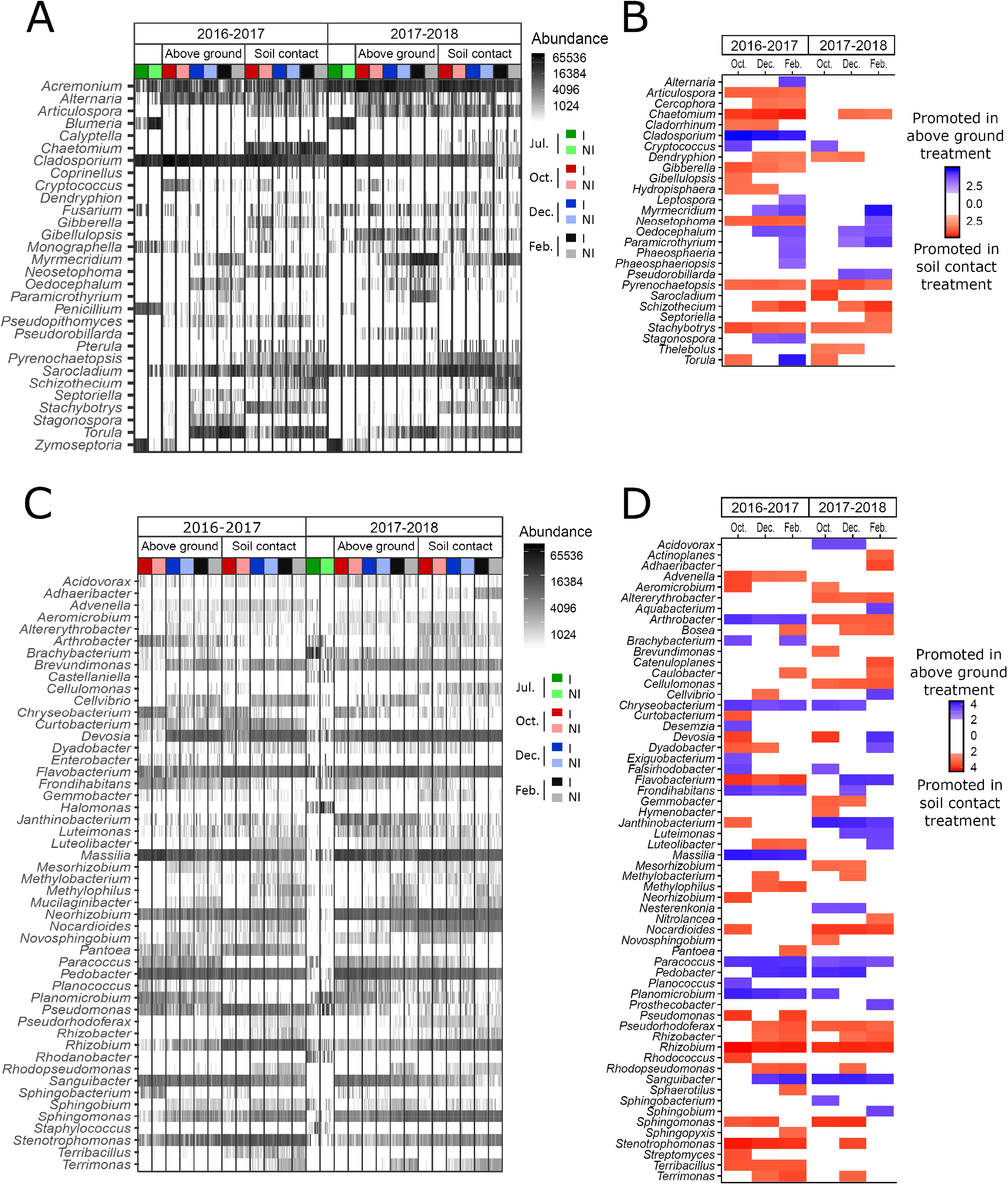
Changes in the relative abundance of microbial taxa over time. (A, C) Diversity and dominance of the 30 most abundant (30/107) fungal genera (A) and the 50 most abundant (50/189) bacterial genera (B) distributed in all samples distinguishing between the different experimental conditions: i.e. cropping season (2016-2017; 2017-2018), contact with soil (“above ground” and “soil contact” treatments), seasonality (July: green; October: red; December: blue; February: gray), and inoculation with *Zymoseptoria tritici* (inoculated: dark hues; non-inoculated: light hues). (B, D) Significant differences in relative abundance of fungal (B) and bacterial (D) genera between the samples in “soil contact” (red) and “above ground” (blue) samples in linear discriminant analysis (LDA). The *Z. tritici* inoculation condition was used as a subclass to avoid interference in the LDA. Only genera with a *p*-value < 0.05 for the Kruskal-Wallis test and an LDA score > 2 are displayed.

Some classes of taxa (e.g. *Bacilli, Sphingobacteria, Betaproteobacteria, Dothideomycetes, Pezizomycetes*) were particularly abundant only in above ground residues, suggesting that they were mostly derived from the plant. By contrast, other classes (e.g. *Alphaproteobacteria, Agaricomycetes, Cytophagia, Gammaproteobacteria*) were more prevalent in residues in contact with soil, suggesting that they originated from the soil (Suppl. Figure 2). The abundance of some classes varied with cropping season (e.g. *Flavobacteria*). Soil contact had a large impact for *Dothideomycetes* and *Bacilli*, which were highly abundant in July, but rapidly decreased in frequency when the residues were in contact with the soil. *Pezizomycetes*, absent in July, colonized only the above ground residues. Conversely, the percentage of reads associated with *Alphaproteobacteria*, which was quite high in July, and *Cytophagia*, which was low in July, increased over time, particularly in residues in contact with the soil. Similarly, *Agarycomycetes*, which was completely absent in July, colonized only residues in contact with the soil.

At the genus level, 87 (excluding “unclassified”) of the 273 genera (60/190 for bacteria; 27/83 for fungi) identified displayed differences in abundance between above ground residues and residues in contact with the soil, for at least one date (Figure 4). For example, *Bosea, Rhizobium, Nocardioides, Pseudomonas*, and *Sphingomonas* were more abundant in residues in contact with the soil, whereas *Cladosporium, Massilia, Paracoccus, Stagonospora* and *Cryptococcus* were more abundant in above ground residues.

### Impact of *Z. tritici* inoculation on microbial communities

The influence of *Z. tritici* inoculation on the RA of residue microbiome members was assessed, through LDA scores. In total, the RA of 115 ASVs (74 bacterial ASVs and 41 fungal ASVs) was significantly affected by *Z. tritici* inoculation, for at least one sampling date (listed in Suppl. Figure 3). The effect of inoculation on microbial communities persisted throughout the experiment, despite the absence of *Zymoseptoria* detection from December onwards (Figure 2). ASVs with significant differences in RA decreased over time for residues in contact with the soil (Suppl. Table 1). By contrast, for above ground residues, the number of differential ASVs increased until December, in both cropping seasons (20 ASVs in December 2016-2017; 31 ASVs in December 2017-2018).

Inoculation with *Z. tritici* decreased the RA of fungal ASVs, including those affiliated to *Sarocladium, Gibellulopsis* and *Blumeria*, and increased the RA of bacterial ASVs affiliated to *Curtobacterium* and *Brachybacterium* (listed in Suppl. Figure 3). The ASVs affected by inoculation differed between above ground residues and residues in contact with soil. The pattern of change (i.e. promoted or inhibited by inoculation) was always the same within a given year, regardless of soil contact conditions. For example, *Brachybacterium* and *Curtobacterium* were promoted by inoculation, in both soil contact conditions, whereas *Sarocladium* was inhibited by inoculation, in both soil contact conditions.

### Impact of the actual presence of *Z. tritici* on microbial communities

Ecological network analyses (ENA) combining bacterial and fungal datasets were performed to predict the potential interactions between *Z. tritici* and members of microbial communities associated with wheat residues.

#### Dynamics of ecological interaction networks

The dataset was split according to the effects previously described (cropping season, seasonality, soil contact conditions). Six ENA were performed per experimental year, corresponding to residue samples in contact with the soil and above ground residues, collected in October, December, and February (Figure 5). The networks for July are presented in Suppl. Figure 4. The mean number of nodes in the network (205.3±47.5) increased over the season (Suppl. Table 1). Overall, networks were sparse, with a mean node degree of 2.76±0.43. For each network, the positive/negative edge ratio decreased over time, reaching 1.0-1.5 in February. Most nodes were common to October, December and February. *Zymoseptoria* ASV was one of the fungal ASV with the largest number of degrees and greatest betweenness (measurement of centrality in a graph based on the shortest paths) for above ground samples in October. By contrast, for samples in contact with soil, it was absent the first year and had low betweenness and degree values for the second year (Figure 6).

**Figure 5.**
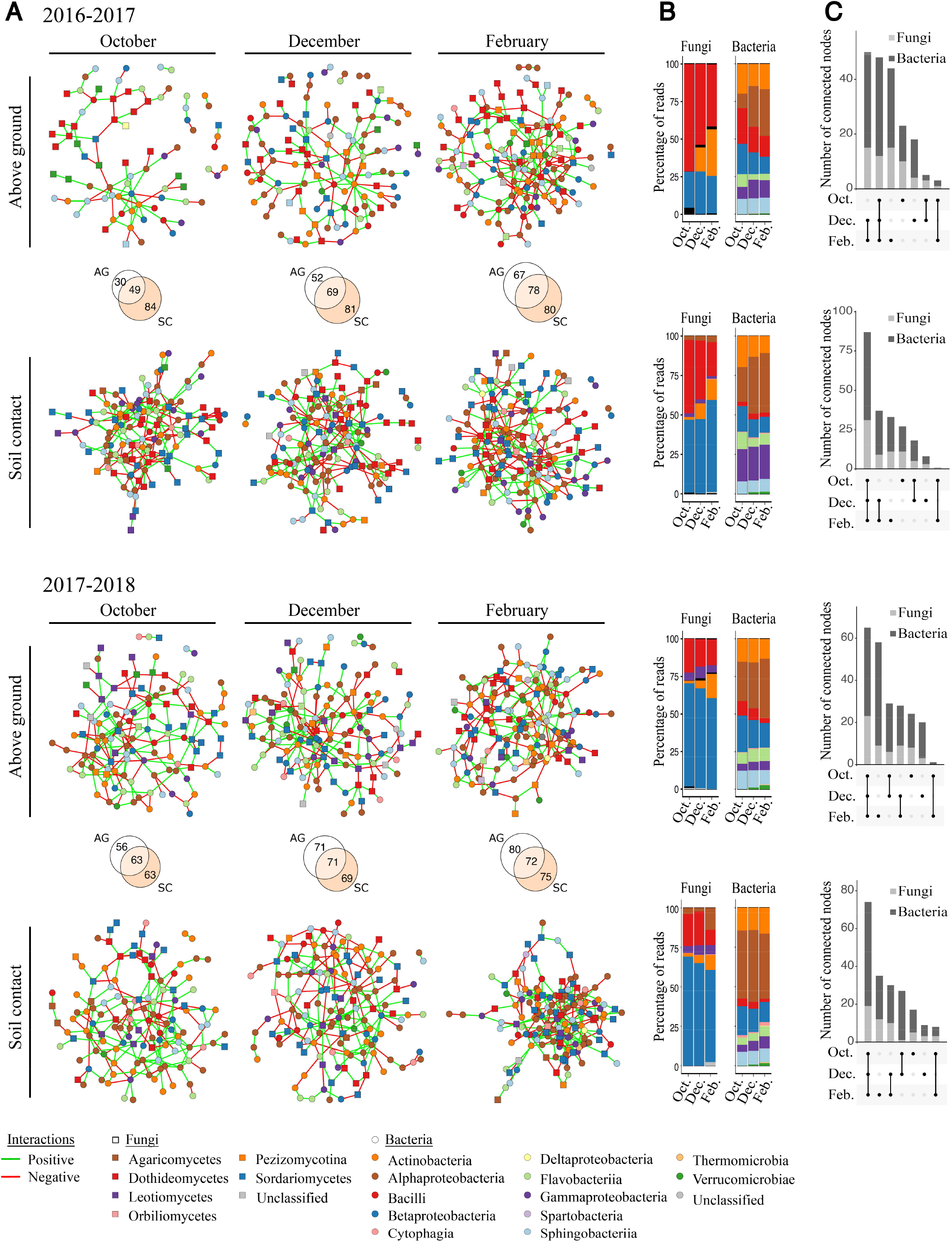
Temporal dynamics of interaction networks. (A) Networks based on bacterial and fungal ASVs combined. In all networks, circles and squares correspond to bacterial and fungal ASVs, respectively, with colors representing class. Isolated nodes are not shown. Edges represent positive (green) or negative (red) interactions. The Venn diagram highlights the number of non-isolated nodes common and specific to “above ground” (AG) and “soil contact” (SC) treatments for each sampling date (October, December, February). (B) Percentage of reads associated with fungal and bacterial classes for each network. Isolated nodes are included. Colors are the same as in (A). (C) Upset plot of bacterial and fungal non-isolated nodes common and specific to sampling date for each treatment.

**Figure 6.**
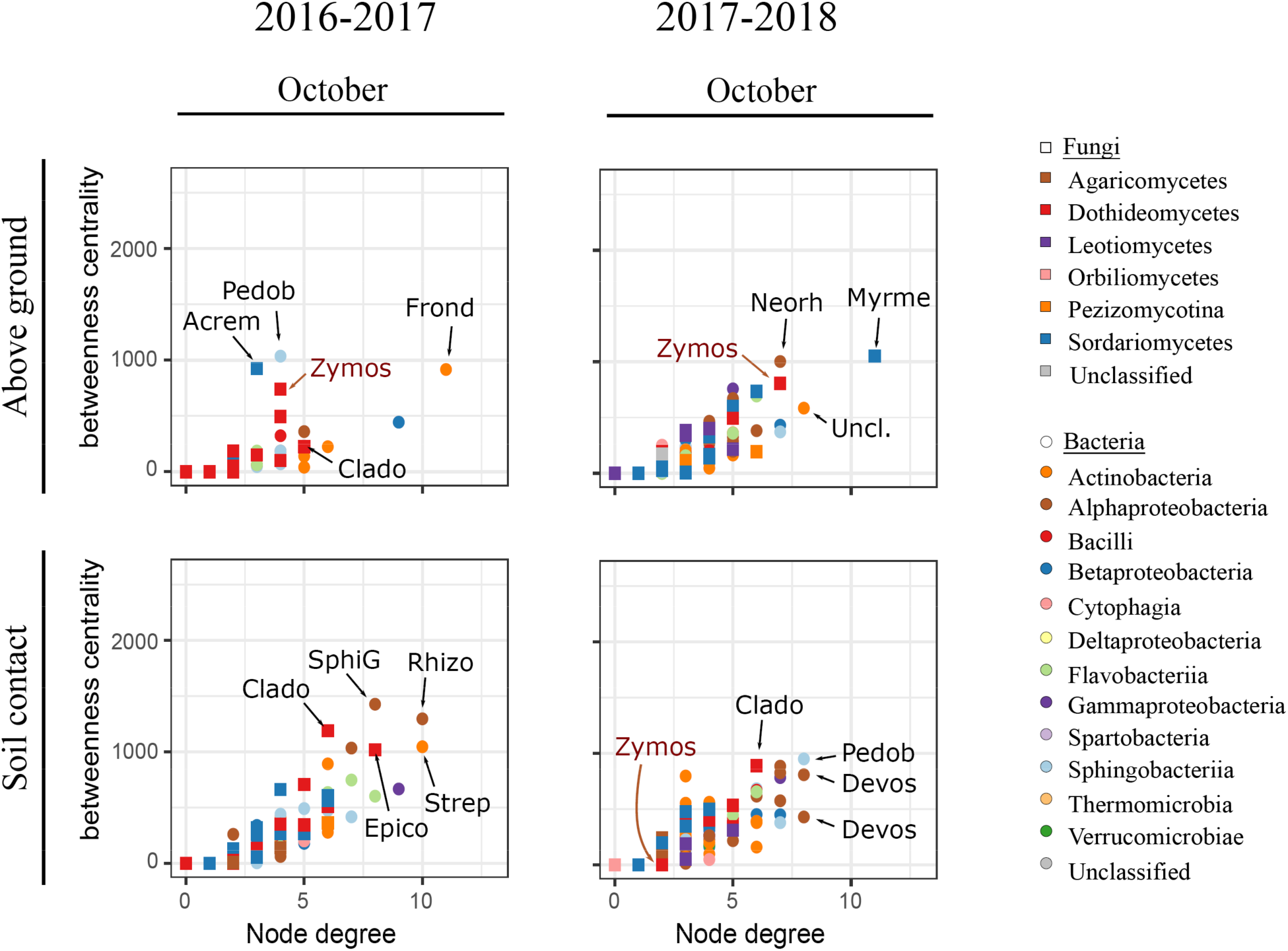
Betweenness centrality and degree of each ASV in the October networks. Nodes with high betweenness centrality and high degree values are considered to be keystone taxa in the networks. The genera of the fungal and bacterial ASVs with the highest degree and centrality are shown: *Acrem(onium)*; *Clado(sporium)*; *Devos(ia)*; *Epico(ccum)*; *Frond(ihabitans)*; *Myrme(cridium)*; *Neorh(izobium)*; *Pedob(acter)*; *Rhizo(bium)*; *SphiG(*=*Sphingomonas)*; *Strep(tomyces)*; *Uncl.(assified)*; *Zymos(eptoria)*. The relationship between betweenness centrality and degree of each ASV in the networks for the other sampling dates (July, December, and February), characterized by a linear regression, are presented in Supplementary Figure 5.

#### *Subnetworks highlighting direct interactions between* Z. tritici *and other microorganisms*

ENA were combined with LDA to investigate the interactions between *Z. tritici* and members of the microbial communities of residues (Figure 7). Only 13 of the 115 ASVs affected by inoculation (LDA) were in direct interaction with Zymoseptoria ASV, indicating an indirect effect of *Z. tritici* on the community (no direct connection between the microorganisms).

**Figure 7.**
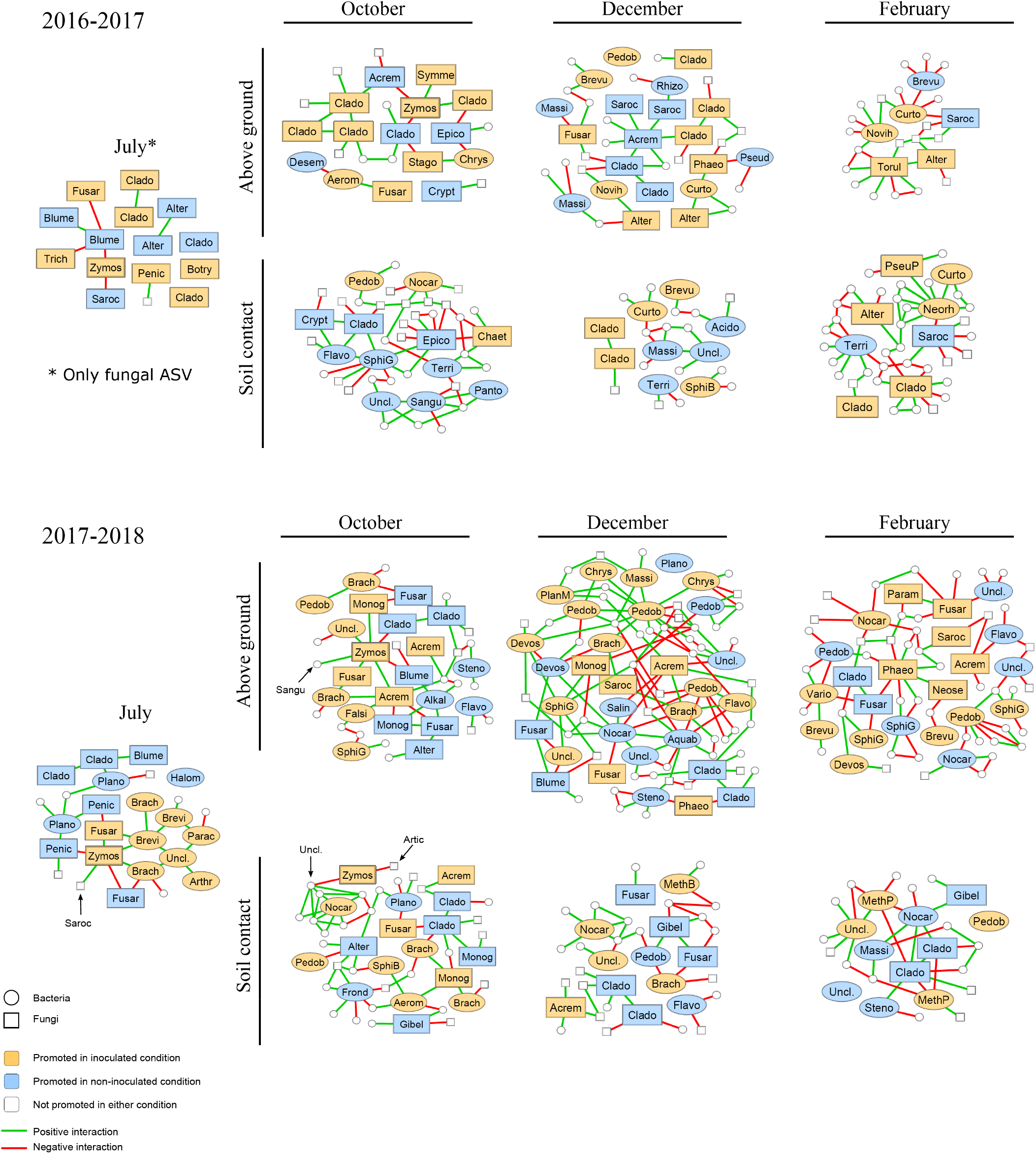
Subnetworks based on the data in Figure 4A and composed of differential bacterial and fungal ASVs identified in residue samples (originating from wheat plants inoculated and non-inoculated with *Zymoseptoria tritici*) and of the first adjacent nodes. Node color corresponds to the results of LefSe differential analysis between inoculated (orange) and non-inoculated (blue) treatments. Only genera with p-values < 0.01 for the Kruskal-Wallis tests and LDA scores > 2 were retained for the plot. The first adjacent nodes of each differential ASV are not named, except for ASVs interacting with *Z. tritici*. Edges represent positive (green) or negative (red) interactions. Differential ASVs are plotted with genus name abbreviations: *Acido(vorax); Acrem(onium); Aerom(icrobium); Alkal(ibacterium); Alter(naria); Aquab(acterium); Arthr(obacter); Blume(ria); Botry(osporium); Brach(ybacterium); Brevi(bacterium); Brevu(ndimonas); Chaet(omium); Chrys(eobacterium); Clado(sporium); Crypt(ococcus); Curto(bacterium); Desem(zia); Devos(ia); Epico(ccum); Falsi(rhodobacter); Flavo(bacterium); Frond(ihabitans); Fusar(ium); Gibel(lulopsis); Halom(onas); Massi(lia); MethB(*=*Methylobacterium); MethP(*=*Methylophilus); Monog(raphella); Neorh(izobium); Neose(tophoma); Nocar(dioides); Novih(erbaspirillum); Panto(ea); Parac(occus); Param(icrothyrium); Pedob(acter); Penic(illium); Phaeo(sphaeria); PhaeP(sphaeriopsis); Plano(coccus); PlanM(*=*Planomicrobium); Pseud(omonas); PseuP(*=*Pseudopithomyces); Rhizo(bium); Rhoda(nobacter); Salin(irepens); Sangu(ibacter); Saroc(ladium); SphiB(*=*Sphingobium); SphiG(*=*Sphingomonas); Stago(nospora); Steno(trophomonas); Symme(trospora); Terri(bacillus); Torul(a); Trich(oderma); Uncl.(assified); Vario(vorax); Zymos(eptoria)*.

Microorganisms with the same differential pattern (i.e. “promoted by inoculation” or “promoted in the absence of inoculation”) did not interact negatively with each other in networks. Conversely, microorganisms with opposite differential patterns systematically interacted negatively with each other. These results highlight the consistency of the LDA and ENA approaches.

The subnetworks generated with microorganisms presenting differential relative abundances and their adjacent nodes were strongly connected: each subnetwork consisted of a principal component and, in some cases, smaller components of less than four nodes (Figure 7).

Only a few direct interactions between *Zymoseptoria* and other microorganisms were highlighted by ENA. Some ASVs affiliated to the same genus had opposite interaction trends with *Zymoseptoria*, such as *Fusarium* ASVs in July 2017, or *Cladosporium* ASVs in October 2016, consistent with the findings of LDA analyses. In some cases, the same ASV had different interaction trends at different sampling dates or in different years. This was the case for *Acremonium* ASVs (negative interaction in October 2016, positive interaction in October 2017). Some genera, such as *Blumeria, Sarocladium*, and *Penicillium*, interacted only negatively with *Zymoseptoria*. *Symmetrospora, Brachybacterium*, and *Monographella* interacted only positively with *Zymoseptoria*.

## Discussion

By sequencing the microbial communities of 420 samples of wheat residues, we obtained a total of 996 bacterial ASVs and 520 fungal ASVs. Using this large dataset, we estimated the potential interactions occurring between a plant pathogen (*Z. tritici*) and the members of microbial communities associated with crop residues in field conditions. By combining two approaches — LDA and ENA — we were able to demonstrate an effect of pathogen infection, even after disappearance of the pathogen, on the structure and composition of the microbial communities during residue degradation.

### Effect of soil contact on microbial communities

Our aim here was not to characterize the organisms colonizing wheat residues, but our findings nevertheless highlight major changes in the microbial community over time for residues in contact with soil. The taxa favored in above ground residues, such as *Cladosporium, Alternaria, Pedobacter* and *Massilia*, were already present on the plant. This is consistent with previous findings showing a decrease in the abundance of these plant-associated taxa during the degradation of residues in contact with soil and the colonization of these residues with soil-borne competitors, such as *Chaetomium, Torula*, and *Nocardioides* [16]. Some fungal genera not present in July were favored by above ground conditions (e.g. *Cryptococcus, Stagonospora*, and *Myrmecridium*). This finding is consistent with our knowledge of fungal dispersal processes, mostly involving aerial spores.

### Decline of *Z. tritici* during residue degradation

*Z. tritici* rapidly decreased to below the limit of detection between October and December. This result is surprising in light of the quantitative epidemiological data acquired for the same plot, which suggested that *Z. tritici* ascospores may be ejected from residues until March [3,37]. The observed decline of *Z. tritici* may be due to lower levels of contamination of adult wheat plants in residues than would be achieved in the field after natural infection. Indeed, in field conditions, *Z. tritici* establishes itself on all parts of the plant (leaves, but also sheaths and stems) through multiple secondary infections, driven by the repeated splash dispersal of asexual spores leading to an accumulation of contaminating raindrops at the points of insertion of the leaf sheaths. The single inoculation event in the greenhouse resulted in contamination principally of the leaves, the organs most exposed to spraying, with relatively little contamination of the stems and sheaths, the parts of the plant most resistant to degradation. Indeed, the results of a previous study [16] support this hypothesis: in the same field, during the same season, *Z. tritici* was detected in wheat residues originating from plants grown in natural conditions until February, and even May, with a similar metabarcoding approach.

### Effect of *Z. tritici* on microbial communities

Endophytes and pathogens induce changes in plant tissues (e.g. necrosis), which may themselves modify the microbial communities inhabiting the plant (e.g. impact of secondary saprophytes or opportunistic pathogens [38]; selection of microorganisms by secondary metabolites produced by microorganisms or the plant [39,40]). This general phenomenon may explain the impact of *Z. tritici* on the microbial communities observed in both LDA and ENA. The impact of *Z. tritici* on residues, even after its decline between October and December, persisted until February, particularly for fungal communities. Within microbial networks, *Z. tritici* was one of the keystone taxa, despite its low abundance, in above ground residues in October (Suppl. Figure 5). The high levels of *Zymoseptoria* in July (between 10 and 40% of reads) account for its central position in the network. The number of microorganisms displaying changes in abundance due to *Z. tritici* inoculation decreased during residue degradation. This finding highlights the resilience of the community (i.e. its ability to return to its original composition after a disturbance, in this case, *Z. tritici* inoculation) [41].

### Specific interactions with *Z. tritici*

Most of the predicted interactions with *Zymoseptoria* involved fungi, such as *Fusarium, Blumeria* or *Cladosporium*. *Z. tritici* infection has been shown to be associated with the accumulation of H_2_O_2_ [42]. This compound is known to inhibit biotroph fungal pathogens [43], such as *Blumeria graminis* [44,45]. This may explain the negative interaction between *Z. tritici* and *B. graminis* in July and October 2017-2018. In addition, *Z. tritici* infection induces leaf necrosis, potentially decreasing wheat susceptibility to *B. graminis*, due to a significant physiological interaction during the latent, endophytic period of *Z. tritici* development [45]. H_2_O_2_ is also known to promote necrotrophic agents, such as *Fusarium*. We detected both positive and negative interactions between *Zymoseptoria* and *Fusarium*, depending on the ASV considered. On adult wheat plants, such differential interactions have been demonstrated in log-linear analyses [46], with both species giving positive results on stem bases and negative results on the upper parts of stems. Positive interactions between *Z. tritici* and *Cladosporium* have also been demonstrated on adult plants [46], consistent with our findings for wheat residues. Although the use of ENA based on bacterial and fungal data sets can introduce many biases (distortion of the microbial community composition due to analysis by separate PCRs, inherent limitations in terms of resolution of the taxonomic markers, etc.), these results lend a biological meaning to the interactions detected, confirming the relevance of network analyses for highlighting ecological interactions within crop residue communities.

*Trichoderma* was more abundant in residues from wheat plants inoculated with *Z. tritici* (July 2016), as shown by LDA (Suppl. Figure 4). Conversely, *Epicoccum* and *Cryptococcus* were more abundant in residues from non-inoculated wheat plants (October 2016). The overabundance of those taxa, described as biocontrol agents in previous studies [34–36,47], was influenced by the presence of the pathogen. However, no direct interactions between *Z. tritici* and these species could be established. This exemplifies the difficulties highlighting beneficial species within complete microbial communities. These difficulties are not specific to the residue compartment and also apply to the spermosphere [48], phyllosphere [49] and rhizosphere compartments [14,50].

### Other interactions

Other interactions between ASVs highlighted in the network analysis were examined in light of published results for fungal pathogens of cereals. For instance, it has already been shown that *B. graminis* growth on barley is inhibited by *Trichoderma harzianum* [51] and Stagonospora norodum [52], that *Stenetrophomas maltophila* attenuates the seedling blight of wheat caused by *F. graminearum* [53], that *Acremonium* zeae has antibiotic activity against *Fusarium verticillioides* [54], and that *Chaetomium* sp. produces compounds (e.g. chaetomin) active against *Alternaria triticimaculans* [34]. Conversely, certain non-pathogenic bacteria were shown to be associated with significantly more disease on wheat caused by *B. graminis* and *Z. tritici* and to “help” Phaeosphaeria nodorum to infect wheat tissues [55]. Newtoon *et al*. [38] has proposed the hypothesis of “induced susceptibility” to explain such an interaction between bacteria and biotroph fungal pathogens.

ENA also suggested that intra-kingdom interactions were favoured over inter-kingdom interactions in certain conditions (Suppl. Table 2). This may reflect differences in ecological niches and dynamics, as illustrated by the temporal changes in microbial communities over a season, with a densification of the networks during residue degradation. Further investigations are required to determine whether inter- or intra-kingdom interactions are more intense, and thus more promising for use in biocontrol engineering. Should we preferentially focus on fungal communities to improve the management of a fungal disease, and on bacterial communities to improve the management of a bacterial disease? The ability to answer this question with the approach developed in this study should be nuanced. Indeed, the weakness associated with separate analysis of fungal and bacterial communities (see above) may have impacted our observation that intra-kingdom interactions were more difficult to discern that inter-kingdom interactions (see below), and may increase the difficulty of identifying actual biological interactions between bacteria and fungi.

### Identification of beneficial species, and potential biocontrol agents

Network models provide new opportunities for enhancing disease management and can be helpful for biocontrol. Our study, combining LDA and ENA based on a metabarcoding approach and differential conditions (plants inoculated with a pathogen or left non-inoculated; plant residues in contact with soil vs. residues not in contact with the soil), fits into the framework described by Poudel *et al*. [56], which considers several types of network analyses, including pathogen-focused analyses, taking into account diseased and healthy plant hosts, with a view to elucidating direct and indirect pathogen-focused interactions within the pathobiome. Network analyses revealed no significant direct interactions between *Z. tritici* and microorganisms reported to be useful biocontrol agents. However, pathogen infection had a strong effect on the entire microbial community present in residues during the course of their degradation. Most of the interactions were difficult to interpret. Several interactions appeared to be transient, changing over time with residue degradation, and their presence or absence depended on whether the residues were in contact with the soil. This suggests that interactions between microorganisms are not stable and can be modified by changes in the environment, for example, or by the arrival of a new microorganism.

Network models, although effective in characterizing putative interactions between ASVs within a microbial community and highlighting changes due to disturbance (e.g. presence of a pathogen, application of fungicides, introduction of a resistance gene in a host plant population, etc.), do not necessarily allow to identify the species concerned by these interactions: indeed, the taxonomic markers employed (16S v4 and ITS1) have inherent limitations in terms of resolution and difficulties for distinguishing microorganisms below the level of genus remain. This is the case for bacteria, but also for a number of fungi, such as those associated with the genus *Alternaria*: some Alternaria sp. are sometimes described as biocontrol agents and others as pathogens, while ITS1 sequences do not allow to distinguish them. Having said that, this type of work combining LDA and ENA based on a metabarcoding approach can be considered as a hypothesis generator or a guide for the targeted isolation of microorganisms that may have the desired biocontrol phenotypes.

The neglect of complex interactions between biocontrol agents and their biotic environment (the plant, the soil and their microbiomes), the physical and chemical properties of which change over time, may account for lower levels of efficacy in field conditions than in laboratory conditions (concerning the phyllosphere, e.g. [38], but also the residue compartment, e.g. [57]). Indeed, several studies have demonstrated the value of studying the effect of entire communities on biotic and abiotic stresses rather than the effects of single species. For example, resistance to *B. cinerea* in *Arabidopsis thaliana* was shown to be not due to a single species, but to the action of the microbiome as a whole [58]. By comparing the structure of microbial communities associated with *Brassica rapa* plants inoculated with the root pathogen *Plasmodiophora brassicae*, Lebreton et al. [14] showed significant shifts in the temporal dynamics of the root and rhizosphere microbiome communities during root infection. Moreover, the rhizospheres of plants infected with *P. brassicae* were significantly more frequently colonized with a *Chytridiomycota* fungus, suggesting interactions between these two microorganisms.

The most frequently studied cases of microbial community effects include “suppressive soils”, which provide defense against soil-borne pathogens, rendering them unable to establish themselves or to persist in the soil or the plant [59]. The basis and dynamics of this disease suppression vary, and suppression may be general or specific, under the control of antibiotic-producing *Pseudomonas* or *Streptomyces* populations, for example [60]. Differences in the composition, structure and diversity of microbial communities on crop residues remain poorly understood, and further studies are required to determine the potential for use in biocontrol not of single agents, but of microbial communities, as for these suppressive soils. Despite this ecological reality, the current perception of biocontrol engineering is still too often limited to the action of a single species, even a single strain, with a direct, strong and durable effect against a plant pathogen.

### Potential utility of the residue microbiome

Improving our understanding of the relationship between biodiversity and ecosystem functioning will require the development of methods integrating microorganisms into the framework of ecological networks. Exhaustive descriptions of microbial diversity combined with ENA are particularly useful for identifying species within microbial communities of potential benefit for disease management [56]. By revealing antagonistic interactions between pathogen species (e.g. *Z. tritici*) and other microorganisms, our study suggests that this strategy could potentially improve the control of residue-borne diseases, as suggested by another recent study on *Fusarium* [17]. This strategy, which has been developed separately for the plant [61,62] and soil [14,50,63] compartments, would undoubtedly benefit from further development on crop residues. Indeed, decreasing the presence of pathogens on residues during the interepidemic period can decrease disease development on subsequent crops [21]. More generally, our case study highlights that an interesting way to use ENA is the definition and comparison of indicators, such as node degree and centrality, to characterize the impact of human-induced perturbations on the microbial component of agroecosystems.

## Conclusion

This study provides one of the first example of research revealing alterations to the crop residue microbiome induced by the presence of a mere residue-borne fungal pathogen using high-throughput DNA sequencing techniques. The strategy developed here can be viewed as a proof-of-concept focusing on crop residues as a particularly rich ecological compartment, with a high diversity of fungal and bacterial taxa originating from both the plant and soil compartments. Our findings pave the way for deeper understanding of the complex interactions between a pathogen, crop residues and other microbial components in the shaping of a plant-protective microbiome, to improve the efficacy of biocontrol agents and to preserve existing beneficial equilibria through the adoption of appropriate agricultural practices.

## Methods

We investigated the effect of *Z. tritici* on the diversity of the wheat microbiome and the effect of the wheat microbiome on *Z. tritici*, by characterizing the composition of the microbial communities of 420 residue samples (210 per year) from plants with and without preliminary *Z. tritici* inoculation. The residues were placed outdoors, either directly in contact with the soil in a field plot or “above ground”, i.e. not in contact with the soil, to assess the effect of their colonization by microorganisms originating from the soil, the plant and the air on the saprophytic development of *Z. tritici*. We investigated the persistence of interactions between the pathogen and the whole microbial community, and changes in those interactions over time, by sampling the residues before exposure to outdoor conditions (in July), and every two months thereafter (in October, December, and February) (Figure 1).

### Preparation of wheat residues

The 420 wheat residue samples were obtained from 60 winter wheat cv. Soissons plants grown in a greenhouse in each of the two years of the study, as described in [64]: two weeks after sowing, seedlings were vernalized for eight weeks in a growth chamber and then transplanted into pots. Three stems per plant were retained. Half the wheat plants were inoculated with a mixture of four *Z. tritici* isolates (two Mat1.1. isolates and two Mat1.2 isolates; [65]) to ensure that sexual reproduction occurred as in natural conditions. This equiproportional conidial suspension was prepared and adjusted to a concentration of 2 × 10^5^ spores.mL^−1^, as previously described [64]. Thirty plants were inoculated at the late heading stage in early May, by spraying with 10 mL of inoculum suspension. The other thirty plants were sprayed with water, as a control. Inoculated and non-inoculated plants were enclosed in transparent plastic bags for three days to ensure moist conditions favoring pathogen infection. Septoria tritici blotch lesions appeared three to four weeks after inoculation (Figure 1A). All plants were kept in the same greenhouse compartment until they reached complete maturity (mid-July).

For each “inoculated” and “non-inoculated” condition, stems and leaves were cut into 2 cm-long pieces and homogenized to generate the “wheat residues”, which were then distributed in 105 nylon bags (1.4 g per bag; Figure 1B) for each set of inoculation conditions, in each year.

### Exposure of residues to natural conditions

Ninety nylon bags were deposited in contact with the soil in a field plot (the “soil contact” treatment) or without contact with the soil (“above ground” residue treatment). Thirty batches of residues (15 inoculated and 15 non-inoculated) were used to characterize the communities present in July before the exposure of the residues in the nylon bags to natural conditions. The field plot (“OWO” in [16]; Grignon experimental station, Yvelines, France; 48°51′N, 1°58′E) was the same in both cropping seasons. It was sown with wheat in 2015-2016, with oilseed rape in 2016-2017, and with wheat in 2017-2018. The 90 bags for the “soil contact” treatment were deposited in the OWO field plot (Figure 1C) in late July, at 15 sampling points 20 m apart (three “inoculated” and three “non-inoculated” bags at each sampling point). The 90 bags of the “above ground” treatment were placed on plastic grids exposed to outdoor conditions and located about 300 m from the OWO field plot (Figure 1D).

We assessed the impact of seasonality on the fungal and bacterial communities on residues by collecting samples of each “inoculated” and “non-inoculated” treatment at three dates (October, December and February): 15 bags from plastic grids (“above ground” treatment) and one bag from each sampling point in the field (“soil contact” treatment) At each date, nylon bags were opened, the residues were rinsed with water and air-dried in laboratory conditions. Residues were then crushed with a Retsch™ Mixer Mill MM 400 for 60 seconds at 30Hz with liquid nitrogen in a Zirconium oxide blender.

### Total DNA extraction

Total DNA was extracted with the DNeasy Plant Mini kit (Qiagen, France), with a slightly modified version of the protocol recommended by the manufacturer. Powdered residues (20 mg), 450 μL of Buffer AP1 preheated to 60°C, RNase A and Reagent DX (450: 1: 1) were mixed vigorously for 15 s in a 2 mL Eppendorf tube. Buffer P3 (130 μL) was added to each tube, which was then shaken manually for 15 s, incubated at −20°C, and centrifuged (1 min, 5000 g). The supernatant (450 μL) was transferred to a spin column and centrifuged (2 min, 20000 g). The filtrate (200 μL) was transferred to a new tube, to which sodium acetate (200 μL, 3 M, pH 5) and cold 2-propanol (600 μL) were added. DNA was precipitated by incubation at −20°C for 30 min and recovered by centrifugation (20 min, 13000 g). The pellet was washed with cold ethanol (70%), dried, and dissolved in 50 μL of AE buffer.

### PCR and Illumina sequencing

Fungal and bacterial communities profiles were analyzed by amplifying ITS1 and the v4 region of the 16S rRNA gene, respectively. Amplifications were performed with ITS1F/ITS2 [66] and 515f/806r [67] primers. All PCRs were run in a total volume of 50 μL, with 1x Qiagen Type-it Multiplex PCR Master Mix (Type-it® Microsatellite PCR kit Cat No./ID: 206243), 0.2 μM of each primer, 1x Q-solution® and 1 μl DNA (approximately 100 ng). The PCR mixture was heated at 95°C for 5 minutes and then subjected to 35 cycles of amplification [95°C (1 min), 60°C (1 min 30 s), 72°C (1 min)] and a final extension step at 72°C (10 min). PCR products were purified with Agencourt® AMPure® XP (Agencourt Bioscience Corp., Beverly, MA). A second round of amplification was performed with 5 μl of purified amplicons and primers containing Illumina adapters and indices. PCR mixtures were heated at 94°C for 1 min, and then subjected to 12 cycles of amplification [94°C (1 min), 55°C (1 min), 68°C (1 min)] and a final extension step at 68°C (10 min). PCR products were purified and quantified with Invitrogen QuantIT™ PicoGreen®. Purified amplicons were pooled in equimolar concentrations, and the final concentration of the library was determined with the qPCR NGS library quantification kit (Agilent). Libraries were sequenced in four independent runs with MiSeq reagent kit v3 (600 cycles).

### Sequence processing

Runs were analyzed separately. Primer sequences were first cut off in the fastq files with Cutadapt [68]. Files were then processed with DADA2 v.1.8.0 [69] according to the recommendations for the “DADA2 Pipeline Tutorial (1.8)” workflow [70], with quality trimming adapted for each run (Suppl. Table 3).

A mock sample consisting of equimolar amounts of DNA from known microorganisms was included in each run (see Suppl. Figure 6) to establish a detection threshold for spurious haplotypes. At a threshold of ≤ 0.3 ‰ of the size of the library, amplicon sequence variants (ASVs) were considered spurious and were removed from the sample. We used the naive Bayesian classifier on RDP trainset 14 [71] and the UNITE 7.1 database [72] to assign ASVs. ASVs assigned to chloroplasts (for bacteria) or unclassified at the phylum level (for bacteria and fungi) were also removed from each sample. Due to the larger proportion of chloroplast sequences among the 16S rRNA gene products obtained from living plant tissues compared to dead tissues, all samples from July 2017 were removed from the analysis.

### Differential community analysis

For microbial community analyses, the total library size of each sample was standardized by normalization by proportion. The experimental conditions taken into account were cropping season (2016-2017 and 2017-2018), seasonality (four sampling dates: July, October, December, and February), inoculation with *Z. tritici* (inoculated and non-inoculated), soil contact (soil contact and above ground treatments). The Shannon diversity index was used to assess the effect of each set of conditions on fungal and bacterial diversity. The divergence of microbial communities between samples was assessed by calculating the Bray-Curtis dissimilarity matrix with the phyloseq package (v 1.24.2 [73]), and then illustrated by MDS and clustering based on the average linkage method (ape package v 5.2. [74]). PERMANOVA was performed with the “margin” option, to test the effect of each factor on communities (adonis2 function, vegan package [75]). Since the July samples were derived from living plant tissues (greenhouse), we carried out a PERMANOVA to test the effects of inoculation (for fungi and bacteria) and season (for fungi only; Table 1), and a PERMANOVA for the other sampling dates together to test the effects of inoculation, season and contact with soil.

A linear discriminant analysis (LDA) implemented in Galaxy [76] (LefSe, http://huttenhower.org/galaxy) was used to characterize the differential abundances of fungal and bacterial taxa between each soil contact condition and each *Z. tritici* inoculation condition. In this analysis, differences in the relative abundance of taxa between treatments were evaluated with a Kruskal-Wallis test; a Wilcoxon test was used to check, by pairwise comparisons, whether all subclasses agreed with the trend identified in the Kruskal-Wallis test. The results were used to construct an LDA model, to discriminate between taxa in the different conditions. For the comparison between “soil contact” and “above ground” treatments, inoculation condition was used as a subclass, with the Wilcoxon test alpha value set at 0.05, and the alpha value of the Kruskal-Wallis test set at 0.01. For the comparison between “inoculated” and “non-inoculated” treatments, the alpha value of the Kruskal Wallis test was set at 0.01 (no subclasses). For both analyses, the threshold for the LDA analysis score was set at 2.0.

### Ecological interaction network analyses

For characterization of interactions within the different wheat residue microbial communities, we performed ecological network analyses (ENA) with SPIEC-EASI [77] for combined bacterial and fungal datasets [78]. The same parameters were used for all networks. The non-normalized abundance dataset was split on the basis of sampling date and soil contact condition. Each of the datasets included 15 inoculated and 15 non-inoculated samples. This choice was based on the following considerations: (i) differences in relative abundance of *Z. tritici* were thus maximal in each dataset; (ii) the variability between samples induced by inoculation was shown to be relatively lower comparatively to sampling date and soil contact condition; (iii) loss in specificity in networks was established to occur because networks are unable to distinguish whether a statistically significant co-occurrence is due to an interaction or rather to a shared habitat preference [79]; (iv) the specificity of networks was established to increase with an increasing number of samples until it plateaued at about 25 [79]. Infrequent ASVs were filtered out by defining a threshold of a minimum of six occurrences, to increase sensitivity of the ENA [79]. We used the neighborhood selection as graphical inference model (Meinshausen and Bühlmann MB method) with SPIEC-EASI, as this method has been shown to outperform most of the other available methods (e.g. CCREPE, SPARCC, SPIEC-EASI (glasso)) [77]. The StARS variability threshold was set at 0.05. Networks were then analyzed with the igraph package (version 1.2.2. [80]). Scripts for network construction and analysis are available from GitHub (see Availability of data and materials).

### Subnetworks for analysis of the *Z. tritici* pathobiome

We used a dual approach to characterize interactions between *Z. tritici* and the other taxa, based on: (i) the LDA scores obtained in differential analyses between *Z. tritici* inoculation conditions (“inoculated” and “non-inoculated” treatments); (ii) ecological network analysis. LDA identified taxa affected by inoculation conditions (definition of classes for samples) and network analysis identified interactions at the sample scale (without prior assumptions). Subnetworks of differential ASVs and their adjacent nodes were established by combining these two approaches. Subnetworks were visualized with Cytoscape V. 3.6.1 [81]

## Declarations

### Acknowledgments

This study was performed in collaboration with the GeT core facility, Toulouse, France (http://get.genotoul.fr/), supported by *France Génomique National Infrastructure*, funded as part of the *“Investissement d’avenir”* program managed by *Agence Nationale pour la Recherche* (contract ANR-10-INBS-09). We thank Angelique Gautier (INRA BIOGER) for preparing the mocks, Sandrine Gelisse (INRA BIOGER), Nathalie Retout (INRA BIOGER) and Christophe Montagnier (INRA Experimental Unit, Thiverval-Grignon) for technical assistance, Anne-Lise Boixel (INRA BIOGER) for assistance with statistical analyses, and Marie-Hélène Balesdent (INRA, UMR BIOGER) for her comments and suggestions to improve the manuscript. We thank Julie Sappa for her help correcting our English.

### Availability of data and materials

The raw sequencing data are available from the European Nucleotide Archive under study accession number PRJEB31818. We provide the command-line script for data analysis and all necessary input files via GitHub (https://github.com/LydieKerdraon/MicrobialNetworkAnalysis-WheatResidues).

### Authors’ contributions

LK, FS, VL, MB conceived the study, participated in its design, and wrote the manuscript. LK conducted the experiments and analyzed the data. FS and VL supervised the project. All authors read and approved the final manuscript.

### Ethics approval and consent to participate

Not applicable

### Consent for publication

Not applicable

## Additional files

**Supplementary Table 1.** Sequence filtering for each run.

| Run | Primers | Sequence number<br>(paired end) | Sequence quality<br>trimming (F/R) | Selection by<br>sequence length | Quality sequence<br>number after<br>DADA2 analysis |
| --- | --- | --- | --- | --- | --- |
| #1 | ITS1F / ITS2 | 10 536 086 (×2) | 220 / 210 | - | 7 164 826 |
| #2 | 515f / 806r | 8 368 872 (×2) | 230 / 200 | 253 bp | 5 562 335 |
| #3 | ITS1F / ITS2 | 10 216 508 (×2) | 220 / 190 | - | 6 965 664 |
| #4 | 515f / 806r | 9 975 344 (×2) | 220 / 170 | 253 bp | 5 734 825 |

**Supplementary Table 2.** Analysis of the proportion of intra-kingdom interactions (between two fungal ASVs and between two bacterial ASVs) and inter-kingdom interactions (between a fungal ASV and a bacterial ASV) in the ecological networks. The statistical significance of the under- or over-representation of inter-kingdom interactions (when F-B residuals < 1 or > 1, respectively) was established by a χ^2^ test of independence performed on the contingency table (χ^2^< 0.001).

| Networks <sup>1</sup> | Number of species |  | Number of interactions |  |  | Theoretical maximum number of interactions |  |  | Residuals |  |  |
| --- | --- | --- | --- | --- | --- | --- | --- | --- | --- | --- | --- |
|  | F <sup>2</sup> | B <sup>3</sup> | F-F | B-B | F-B | F-F <sup>4</sup> | B-B <sup>5</sup> | F-B <sup>6</sup> | F-F | B-B | F-B |
| Oct. 2016-2017, above ground | 32 | 73 | 17 | 57 | 17 | 496 | 2628 | 2336 | 1.856 | 0.545 | -1.939 |
| Oct. 2016-2017, contact with soil | 52 | 90 | 32 | 121 | 69 | 1326 | 4005 | 4680 | 1.060 | -0.736 | 0.358 |
| Dec. 2016-2017, above ground | 39 | 100 | 19 | 86 | 42 | 741 | 4950 | 3900 | 0.340 | 0.036 | -0.266 |
| Dec. 2016-2017, contact with soil | 51 | 105 | 25 | 145 | 74 | 1275 | 5460 | 5355 | -0.772 | 0.236 | 0.160 |
| Feb. 2016-2017, above ground | 48 | 107 | 30 | 110 | 65 | 1128 | 5671 | 5136 | 1.109 | -0.866 | 0.508 |
| Feb. 2016-2017, contact with soil | 55 | 110 | 32 | 136 | 90 | 1485 | 5995 | 6050 | 0.208 | -1.170 | 1.505 |
| July 2017-2018 | 16 | 60 | 8 | 45 | 9 | 120 | 1770 | 960 | 0.216 | 1.476 | -2.201 |
| Oct. 2017-2018, above ground | 39 | 93 | 28 | 93 | 62 | 741 | 4278 | 3627 | 1.309 | -1.321 | 1.019 |
| Oct. 2017-2018, contact with soil | 31 | 109 | 17 | 140 | 43 | 465 | 5886 | 3379 | -1.413 | 2.172 | -2.144 |
| Dec. 2017-2018, above ground | 42 | 110 | 22 | 119 | 68 | 861 | 5995 | 4620 | -0.598 | -0.253 | 0.733 |
| Dec. 2017-2018, contact with soil | 35 | 116 | 19 | 149 | 50 | 595 | 6670 | 4060 | -1.384 | 1.949 | -1.849 |
| Feb. 2017-2018, above ground | 44 | 119 | 27 | 136 | 72 | 946 | 7021 | 5236 | -0.207 | -0.081 | 0.244 |
| Feb. 2017-2018, contact with soil | 46 | 114 | 26 | 135 | 91 | 1035 | 6441 | 5244 | -0.752 | -0.978 | 1.845 |

<sup>1</sup> according to sampling date, cropping season and contact with soil
<sup>2</sup> fungal species
<sup>3</sup> bacterial species
<sup>4</sup> estimated by $C_2^{n_F} = \frac{n_F!}{2 \times (n_F-2)!}$
<sup>5</sup> estimated by $C_2^{n_B}$
<sup>6</sup> estimated by $n_F \times n_B$

**Supplementary Table 3.**
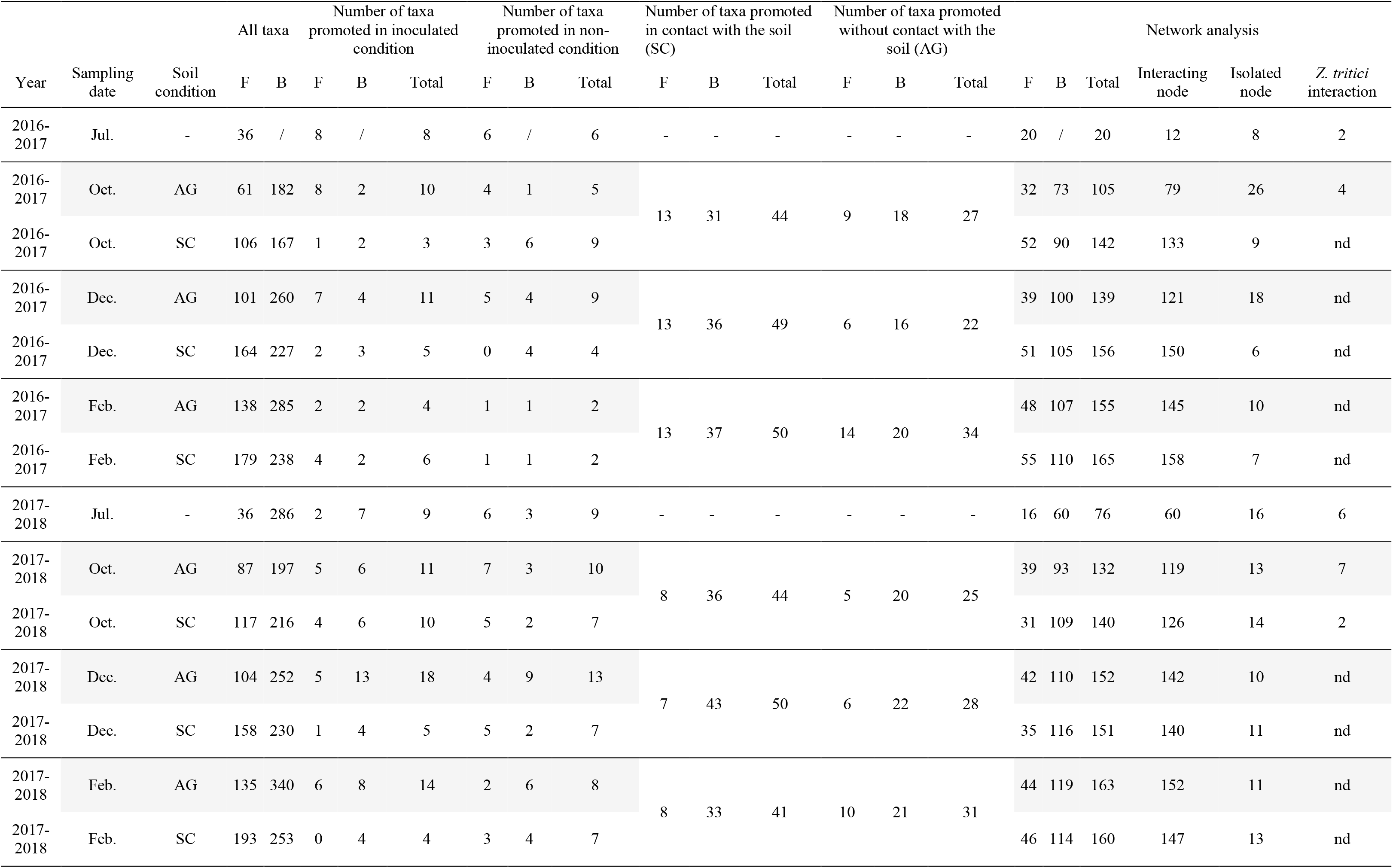
Number of ASVs detected for each analysis performed on the dataset and properties of residue microbial ecological networks.

**Supplementary Figure 1.**
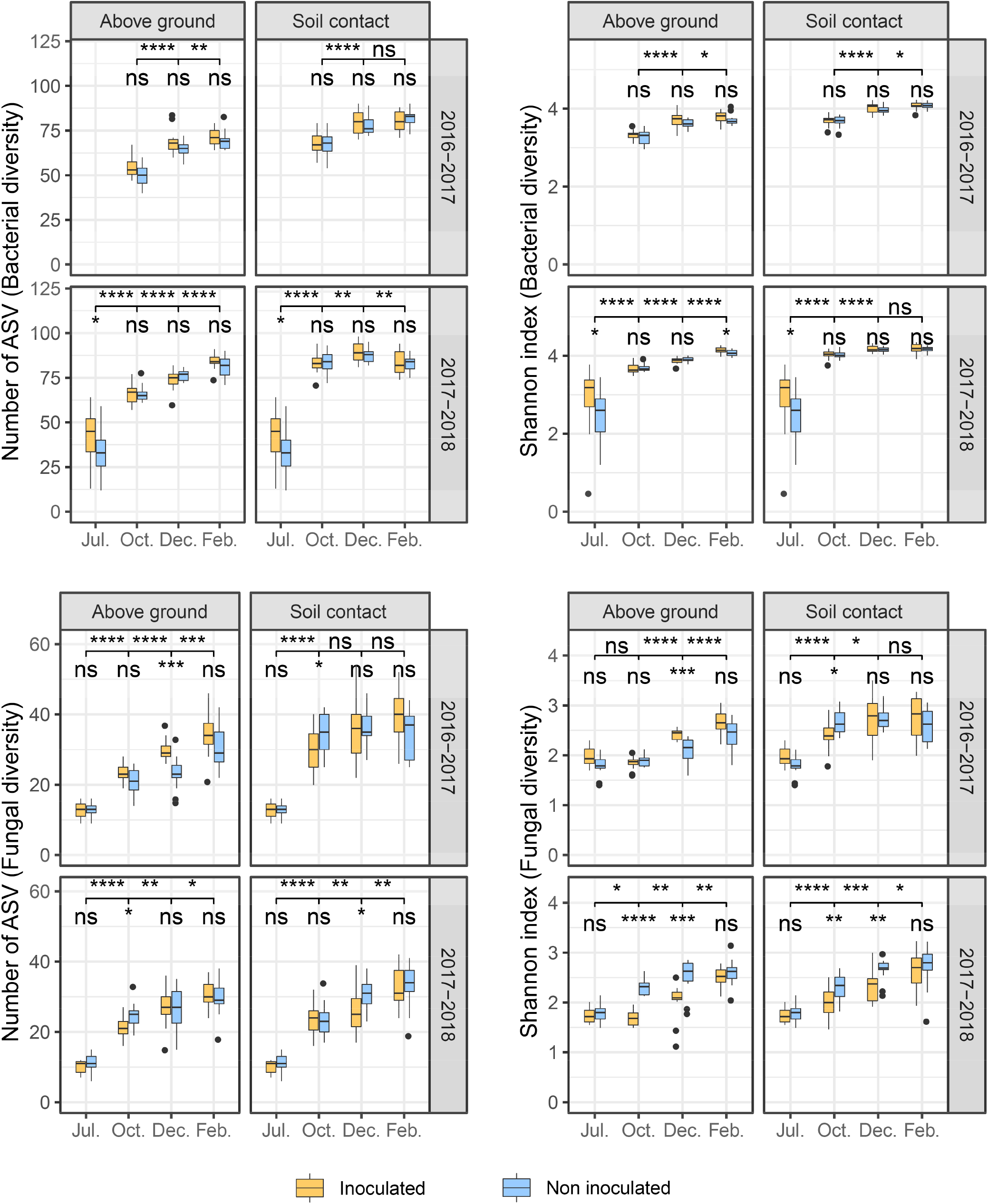
Alpha diversity of microbial communities associated with residues. Observed richness (number of ASVs) and diversity (Shannon index), in four sets of experimental conditions (cropping season, contact with soil, seasonality, *Zymoseptoria tritici* inoculation). Each box represents the distribution of the number of ASVs and Shannon index for 15 sampling points per treatment. Wilcoxon tests were performed for inoculation condition (inoculated, non-inoculated) and sampling date (July, October, December, February). Wilcoxon tests were performed for inoculation condition, and between sampling dates (NS: not significant; * *p*-value < 0.05; ** *p*-value < 0.01; *** *p*-value < 0.001).

**Supplementary Figure 2.**
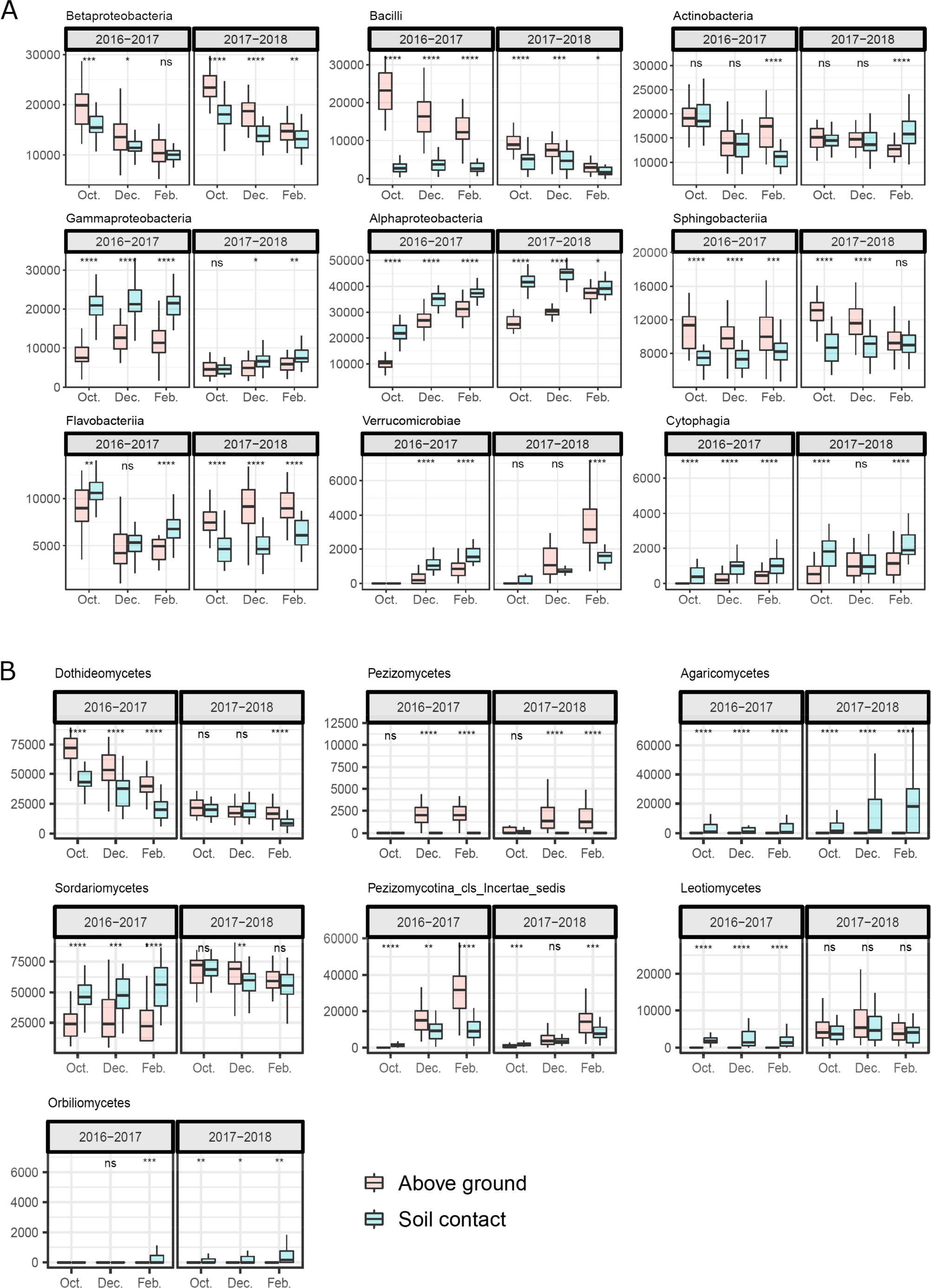
Seasonal shift, from October to February, in the relative abundance of a selection of bacterial (A) and fungal (B) classes present on wheat residues (originating from wheat plants inoculated and not inoculated with *Zymoseptoria tritici*) according to cropping season (2016-2017, 2017-2018) and soil contact condition (in contact with the soil or above ground). Each box represents the distribution of class relative abundances for the 15 sampling points per treatment. Wilcoxon tests were performed for soil contact condition (NS: not significant; * *p*-value < 0.05; ** *p*-value < 0.01; *** *p*-value < 0.001).

**Supplementary Figure 3.**
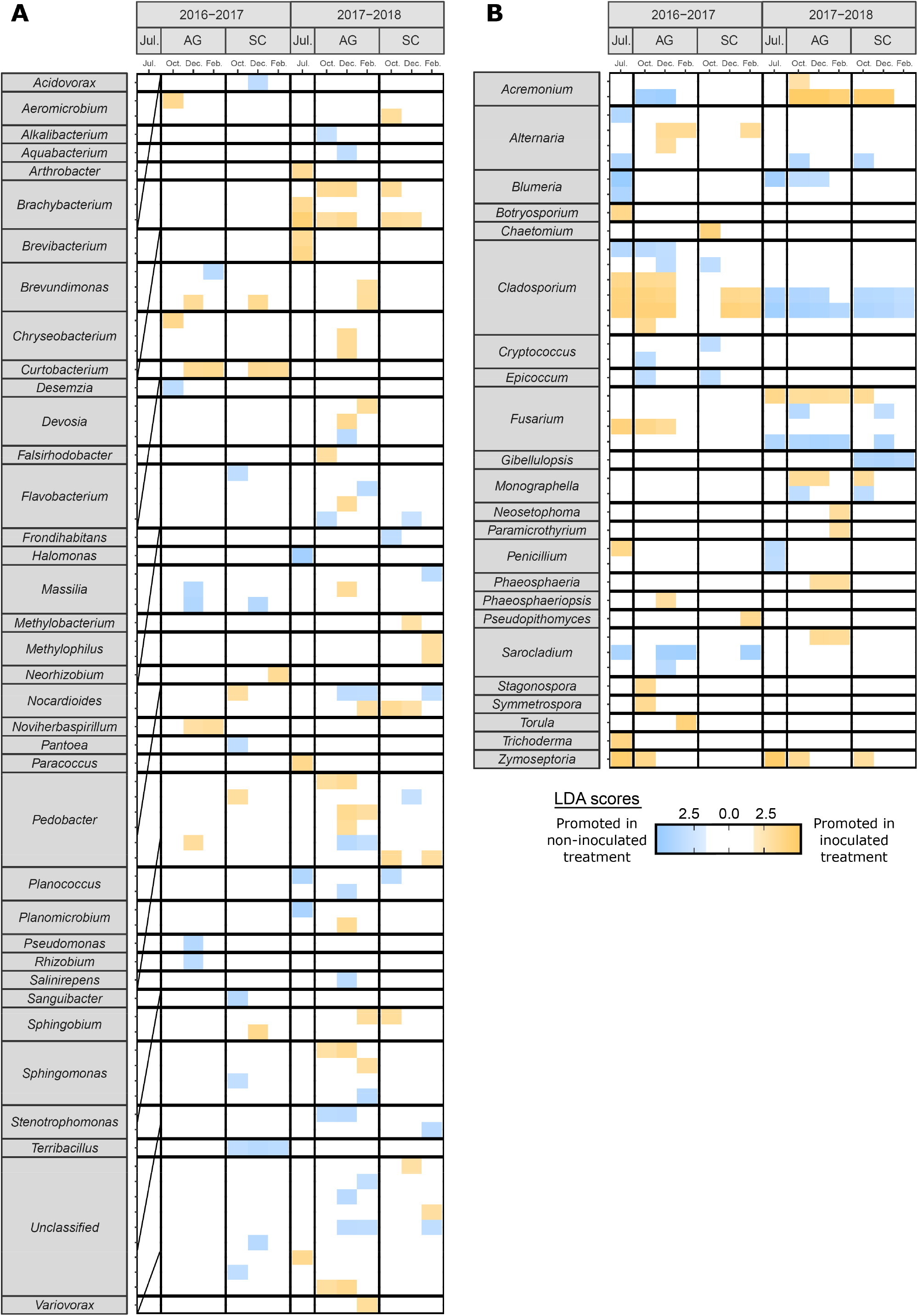
Significant differences in the dominance of fungal and bacterial genera between wheat residues originating from inoculated (orange) and non-inoculated (blue) wheat plants in linear discriminant analyses (LDA), according to three sets of experimental conditions (cropping season, soil contact, seasonality). Only ASVs with *p*-values < 0.01 for the Kruskal-Wallis test and LDA scores > 2 are displayed.

**Supplementary Figure 4.**
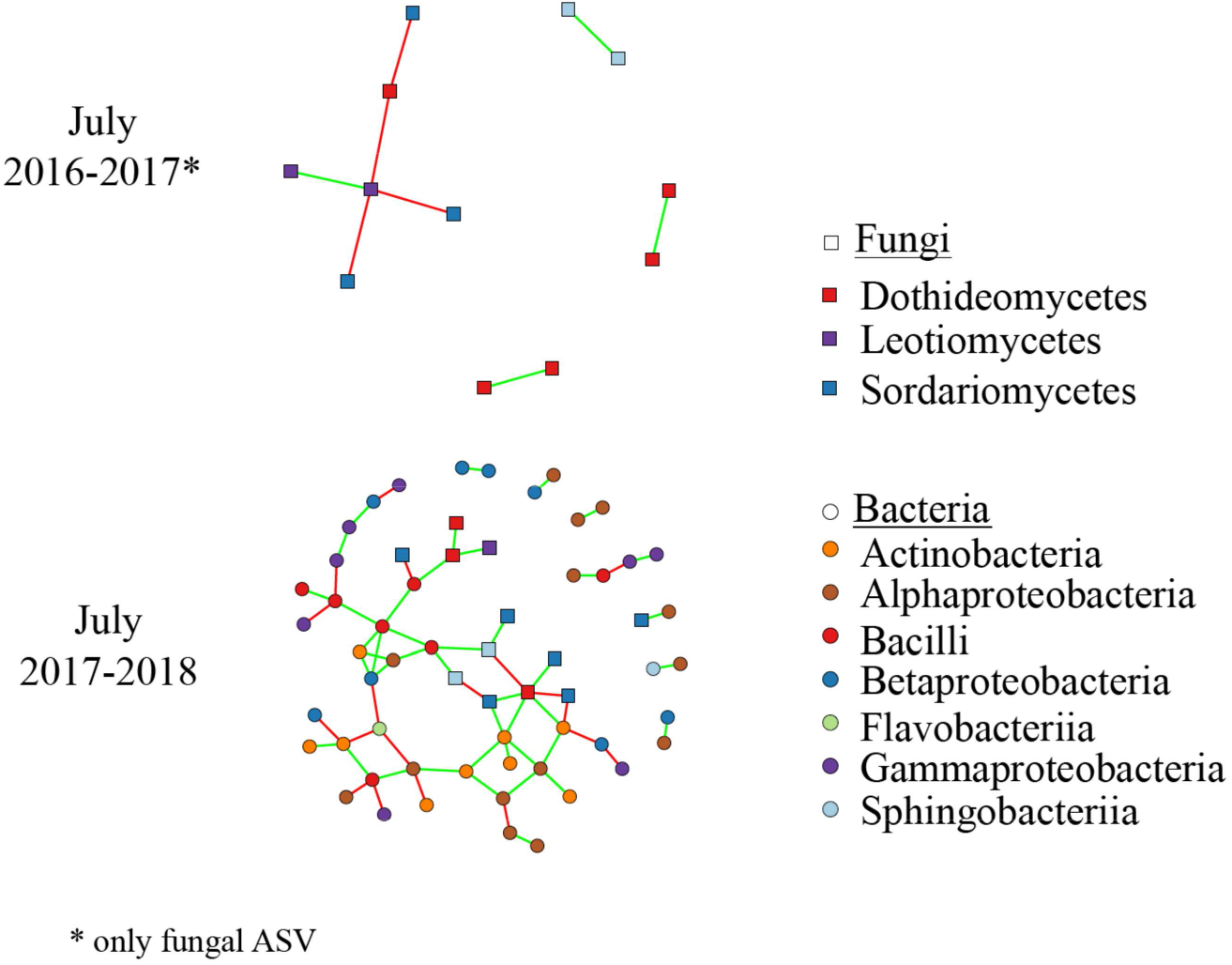
Interaction networks based on bacterial and fungal ASVs combined for July (no contact with soil) for each cropping season (2016-2017, 2017-2018). Circles and squares correspond to bacterial and fungal ASVs, respectively, with colors represent classes. Isolated nodes are not shown. Edges represent positive (green) or negative (red) interactions.

**Supplementary Figure 5.**
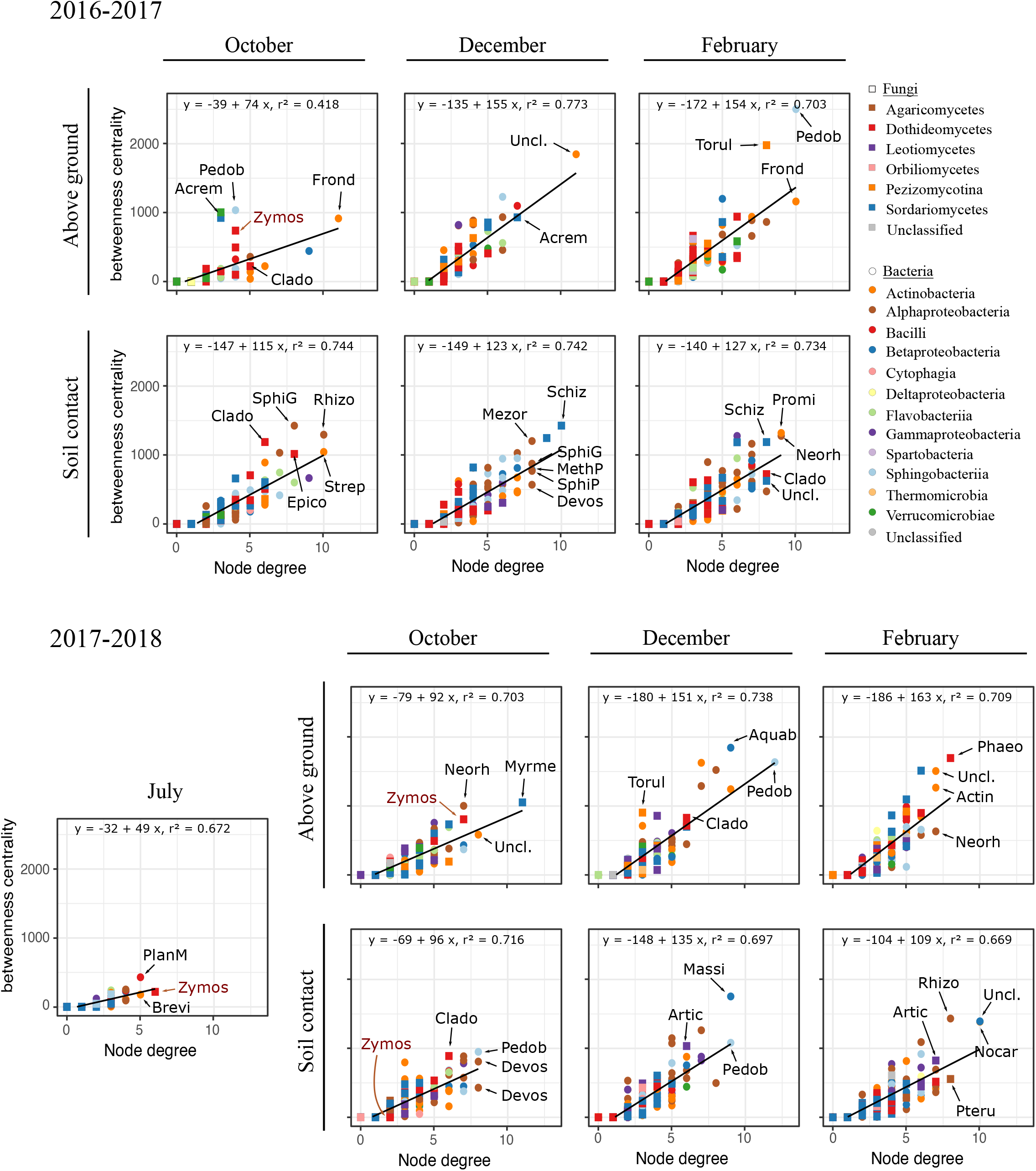
Betweenness, centrality and degree of each ASV in the networks. Nodes with high betweenness, centrality and high degree values are considered to be keystone taxa in the networks. The genera of the fungal and bacterial ASVs with the highest degree and centrality are indicated: *Acrem(onium); Actin(oplanes); Aquab(acterium); Artic(ulospora); Brevi(bacterium); Clado(sporium); Devos(ia); Epico(ccum); Frond(ihabitans); Massi(lia); Mesor(hizobium); MethP(*=*Methylophilus); Myrme(cridium); Neorh(izobium); Nocar(dioides); Pedob(acter); Phaeo(sphaeria); PlanM(*=*Planomicrobium); Promi(cromonospora); Pteru(la); Rhizo(bium); Schiz(othecium); SphiG(*=*Sphingomonas); SphiP(*=*Sphingopyxis); Strep(tomyces); Torul(a); Uncl.(assified); Zymos(eptoria)*.

**Supplementary Figure 6.**
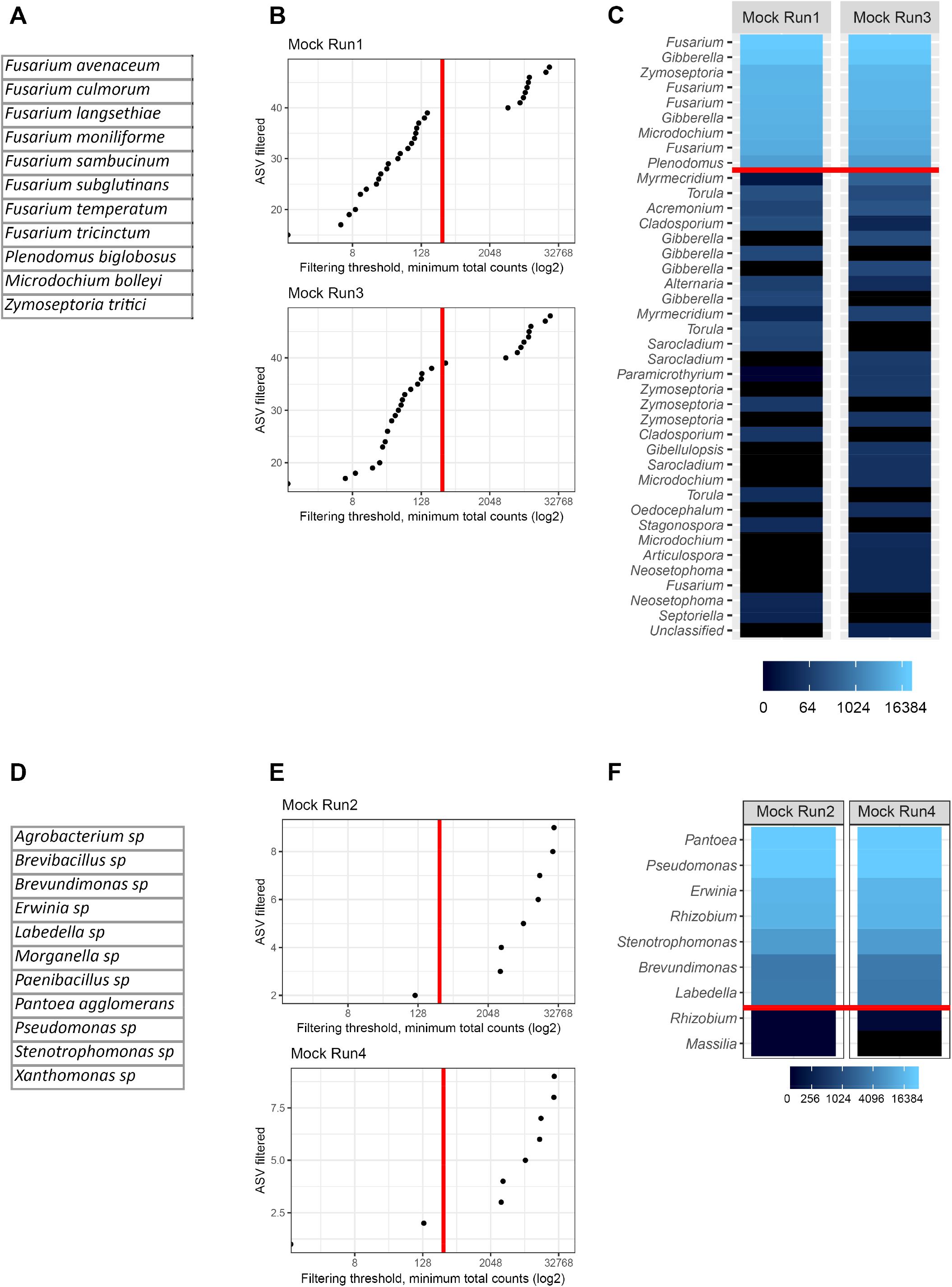
Mocks analysis for the two fungal sequencing runs (A, C) and the two bacterial sequencing runs (D, F). (A, D) Composition of the mocks. All microbial DNAs were pooled at equimolar concentrations. (B, E) Filter on the relative abundance of ASVs. The library size was normalized by proportion before analysis. The red line corresponds to a threshold at 3 ‰ of the size of the library. (C, F) ASVs detected in each mock. The 40 most abundant fungal ASVs are indicated (C), whereas all bacterial ASVs are indicated (F). The name of the ASVs corresponds to the taxonomic affiliation to the genus. All genera present in fungal mocks were detected (*Gibberella* and *Fusarium* are synonymous), while some bacterial genera were not detected in bacterial mocks, which differed only from one ASV. The red line corresponds to a threshold at 3 ‰ of the size of the library.

## References

[1] Fones H, Gurr S. The impact of Septoria tritici blotch disease on wheat: An EU perspective. Fungal Genet Biol 2015;79:3–7.

[2] Quaedvlieg W, Kema G, Groenewald J, Verkley G, Seifbarghi S, Razavi M, et al. *Zymoseptoria* gen. nov.: a new genus to accommodate *Septoria*-like species occurring on graminicolous hosts. Persoonia Mol Phylogeny Evol Fungi 2011;26:57.

[3] Suffert F, Sache I, Lannou C. Early stages of Septoria tritici blotch epidemics of winter wheat: build-up, overseasoning, and release of primary inoculum. Plant Pathol 2011;60:166–77.

[4] Suffert F, Delestre G, Gélisse S. Sexual reproduction in the fungal foliar pathogen *Zymoseptoria tritici* is driven by antagonistic density-dependence mechanisms 2018. Microb Ecol 2018;77:110–23.

[5] Cools HJ, Fraaije BA. Are azole fungicides losing ground against Septoria wheat disease? Resistance mechanisms in *Mycosphaerella graminicola*. Pest Manag Sci 2008;64:681–4.

[6] Estep LK, Zala M, Anderson NP, Sackett KE, Flowers M, McDonald BA, et al. First report of resistance to QoI fungicides in North American populations of *Zymoseptoria tritici*, causal agent of Septoria tritici blotch of wheat. Plant Dis 2013;97:1511.

[7] Estep LK, Torriani SFF, Zala M, Anderson NP, Flowers MD, McDonald BA, et al. Emergence and early evolution of fungicide resistance in North American populations of *Zymoseptoria tritici*. Plant Pathol 2015;64:961–71.

[8] Hayes LE, Sackett KE, Anderson NP, Flowers MD, Mundt CC. Evidence of selection for fungicide resistance in *Zymoseptoria tritici* populations on wheat in western Oregon. Plant Dis 2016;100:483–9.

[9] Cowger C, Hoffer M, Mundt C. Specific adaptation by *Mycosphaerella graminicola* to a resistant wheat cultivar. Plant Pathol 2000;49:445–51.

[10] Berendsen RL, Vismans G, Yu K, Song Y, Jonge R, Burgman WP, et al. Disease-induced assemblage of a plant-beneficial bacterial consortium. ISME J 2018;12:1496.

[11] Toju H, Peay KG, Yamamichi M, Narisawa K, Hiruma K, Naito K, et al. Core microbiomes for sustainable agroecosystems. Nat Plants 2018;4:247–57.

[12] Cho I, Blaser MJ. The Human Microbiome: at the interface of health and disease. Nat Rev Genet 2012;13:260–70.

[13] Jakuschkin B, Fievet V, Schwaller L, Fort T, Robin C, Vacher C. Deciphering the Pathobiome: Intra- and interkingdom interactions involving the pathogen *Erysiphe alphitoides*. Microb Ecol 2016;72:870–80.

[14] Lebreton L, Guillerm-Erckelboudt A-Y, Gazengel K, Linglin J, Ourry M, Glory P, et al. Temporal dynamics of bacterial and fungal communities during the infection of Brassica rapa roots by the protist *Plasmodiophora brassicae*. PloS One 2019;14:e0204195.

[15] Vayssier-Taussat M, Albina E, Citti C, Cosson J-F, Jacques M-A, Lebrun M-H, et al. Shifting the paradigm from pathogens to pathobiome: new concepts in the light of meta-omics. Front Cell Infect Microbiol 2014;4.

[16] Kerdraon L, Balesdent M-H, Barret M, Laval V, Suffert F. Crop residues in wheat-oilseed rape rotation system: a pivotal, shifting platform for microbial meetings. Microb Ecol 2019;77:931–45.

[17] Cobo-Díaz JF, Baroncelli R, Le Floch G, Picot A. Combined metabarcoding and co-occurrence network analysis to profile the bacterial, fungal and *Fusarium* communities and their interactions in maize stalks. Front Microbiol 2019;10:261.

[18] Hadas A, Kautsky L, Goek M, Erman Kara E. Rates of decomposition of plant residues and available nitrogen in soil, related to residue composition through simulation of carbon and nitrogen turnover. Soil Biol Biochem 2004;36:255–66.

[19] Pascault N, Ranjard L, Kaisermann A, Bachar D, Christen R, Terrat S, et al. Stimulation of different functional groups of bacteria by various plant residues as a driver of soil priming effect. Ecosystems 2013;16:810–22.

[20] Pascault N, Cécillon L, Mathieu O, Hénault C, Sarr A, Lévêque J, et al. In situ dynamics of microbial communities during decomposition of wheat, rape, and alfalfa residues. Microb Ecol 2010;60:816–28.

[21] Kerdraon L, Laval V, Suffert F. Microbiomes and pathogen survival in crop residues, an ecotone between plant and soil. Phytobiomes J, accepted.

[22] Nicolardot B, Bouziri L, Bastian F, Ranjard L. A microcosm experiment to evaluate the influence of location and quality of plant residues on residue decomposition and genetic structure of soil microbial communities. Soil Biol Biochem 2007;39:1631–44.

[23] Dugan FM, Lupien SL, Hernandez-Bello M, Peever TL, Chen W. Fungi resident in chickpea debris and their suppression of growth and reproduction of *Didymella rabiei* under laboratory conditions. J Phytopathol 2005;153:431–439.

[24] Fernandez MR. The effect of *Trichoderma harzianum* on fungal pathogens infesting wheat and black oat straw. Soil Biol Biochem 1992;24:1031–4.

[25] Inch S, Gilbert J. Effect of *Trichoderma harzianum* on perithecial production of Gibberella zeae on wheat straw. Biocontrol Sci Technol 2007;17:635–46.

[26] Bujold I, Paulitz TC, Carisse O. Effect of *Microsphaeropsis* sp. on the production of perithecia and ascospores of *Gibberella zeae*. Plant Dis 2001;85:977–84.

[27] Palazzini JM, Groenenboom-de Haas B, Torres AM, Köhl J, Chulze SN. Biocontrol and population dynamics of *Fusarium* spp. on wheat stubble in Argentina. Plant Pathol 2013;62:859–66.

[28] Schöneberg A, Musa T, Voegele R, Vogelgsang S. The potential of antagonistic fungi for control of *Fusarium graminearum* and *Fusarium crookwellense* varies depending on the experimental approach. J Appl Microbiol 2015;118:1165–79.

[29] Palazzini JM, Yerkovich N, Alberione E, Chiotta M, Chulze SN. An integrated dual strategy to control *Fusarium graminearum* sensu stricto by the biocontrol agent *Streptomyces* sp. RC 87B under field conditions. Plant Gene 2017;9:13–18.

[30] Legrand F, Picot A, Cobo-Díaz JF, Chen W, Le Floch G. Challenges facing the biological control strategies for the management of Fusarium Head Blight of cereals caused by *F. graminearum*. Biol Control 2017;113:26–38.

[31] Perez C, Dill-Macky R, Kinkel LL. Management of soil microbial communities to enhance populations of *Fusarium graminearum*-antagonists in soil. Plant Soil 2008;302:53–69.

[32] Kildea S, Ransbotyn V, Khan MR, Fagan B, Leonard G, Mullins E, et al. *Bacillus megaterium* shows potential for the biocontrol of Septoria tritici blotch of wheat. Biol Control 2008;47:37–45.

[33] Levy E, Eyal Z, Chet I. Suppression of Septoria tritici blotch and leaf rust on wheat seedling leaves by *pseudomonads*. Plant Pathol 1988;37:551–7.

[34] Perelló A, Simón MR, Arambarri AM, Cordo CA. Greenhouse screening of the saprophytic resident microflora for control of leaf spots of wheat (*Triticum aestivum*). Phytoparasitica 2001;29:341–351.

[35] Stocco MC, Mónaco CI, Abramoff C, Lampugnani G, Salerno G, Kripelz N, et al. Selection and characterization of Argentine isolates of *Trichoderma harzianum* for effective biocontrol of Septoria leaf blotch of wheat. World J Microbiol Biotechnol 2016;32.

[36] Cordo CA, Monaco CI, Segarra CI, Simon MR, Mansilla AY, Perelló AE, et al. *Trichoderma* spp. as elicitors of wheat plant defense responses against *Septoria tritici*. Biocontrol Sci Technol 2007;17:687–98.

[37] Morais D, Sache I, Suffert F, Laval V. Is the onset of septoria tritici blotch epidemics related to the local pool of ascospores? Plant Pathol 2016;65:250–60.

[38] Newton A, Gravouil C, Fountaine J. Managing the ecology of foliar pathogens: ecological tolerance in crops. Ann Appl Biol 2010;157:343–59.

[39] Hartmann A, Schmid M, Tuinen D van, Berg G. Plant-driven selection of microbes. Plant Soil 2009;321:235–57.

[40] Pusztahelyi T, Holb IJ, Pócsi I. Secondary metabolites in fungus-plant interactions. Front Plant Sci 2015;6:573.

[41] Allison SD, Martiny JBH. Resistance, resilience, and redundancy in microbial communities. Proc Natl Acad Sci 2008;105:11512–9.

[42] Shetty NP, Mehrabi R, Lütken H, Haldrup A, Kema GH, Collinge DB, et al. Role of hydrogen peroxide during the interaction between the hemibiotrophic fungal pathogen *Septoria tritici* and wheat. New Phytol 2007;174:637–47.

[43] Thordal-Christensen H, Zhang Z, Wei Y, Collinge DB. Subcellular localization of H_2_O_2_ in plants. H_2_O_2_ accumulation in papillae and hypersensitive response during the barley— powdery mildew interaction. Plant J 1997;11:1187–94.

[44] Trujillo M, Kogel K-H, Hückelhoven R. Superoxide and hydrogen peroxide play different roles in the nonhost interaction of barley and wheat with inappropriate formae speciales of *Blumeria graminis*. Mol Plant Microbe Interact 2004;17:304–12.

[45] Orton ES, Brown JK. Reduction of growth and reproduction of the biotrophic fungus *Blumeria graminis* in the presence of a necrotrophic pathogen. Front Plant Sci 2016;7:742.

[46] Grudzinska-Sterno M, Yuen J, Stenlid J, Djurle A. Fungal communities in organically grown winter wheat affected by plant organ and development stage. Eur J Plant Pathol 2016;146:401–17.

[47] Perelló AE, Moreno MV, Mónaco C, Simón MR, Cordo C. Biological control of Septoria tritici blotch on wheat by *Trichoderma* spp. under field conditions in Argentina. BioControl 2009;54:113–22.

[48] Nelson EB. Microbial dynamics and interactions in the spermosphere. Annu Rev Phytopathol 2004;42:271–309.

[49] Saleem M, Meckes N, Pervaiz ZH, Traw MB. Microbial interactions in the phyllosphere increase plant performance under herbivore biotic stress. Front Microbiol 2017;8:41.

[50] Whipps JM. Microbial interactions and biocontrol in the rhizosphere. J Exp Bot 2001;52:487–511.

[51] Haugaard H, Lyngs Jørgensen H, Lyngkjær M, Smedegaard-Petersen V, Collinge DB. Control of *Blumeria graminis f. sp. hordei* by treatment with mycelial extracts from cultured fungi. Plant Pathol 2001;50:552–60.

[52] Weber G, Gülec S, Kranz J. Interactions between *Erysiphe graminis* and *Septoria nodorum* on wheat. Plant Pathol 1994;43:158–63.

[53] Dal Bello G, Monaco C, Simon M. Biological control of seedling blight of wheat caused by *Fusarium graminearum* with beneficial rhizosphere microorganisms. World J Microbiol Biotechnol 2002;18:627–36.

[54] Wicklow DT, Poling SM. Antimicrobial activity of pyrrocidines from *Acremonium zeae* against endophytes and pathogens of maize. Phytopathology 2009;99:109–15.

[55] Dewey F, Wong YL, Seery R, Hollins T, Gurr S. Bacteria associated with *Stagonospora* (*Septoria*) *nodorum* increase pathogenicity of the fungus. New Phytol 1999;144:489–97.

[56] Poudel R, Jumpponen A, Schlatter D, Paulitz T, Gardener BM, Kinkel LL, et al. Microbiome networks: a systems framework for identifying candidate microbial assemblages for disease management. Phytopathology 2016;106:1083–96.

[57] Luongo L, Galli M, Corazza L, Meekes E, Haas LD, Van Der Plas CL, et al. Potential of fungal antagonists for biocontrol of *Fusarium spp*. in wheat and maize through competition in crop debris. Biocontrol Sci Technol 2005;15:229–42.

[58] Ritpitakphong U, Falquet L, Vimoltust A, Berger A, Métraux J-P, L’Haridon F. The microbiome of the leaf surface of *Arabidopsis* protects against a fungal pathogen. New Phytol 2016;210:1033–43.

[59] Weller DM, Raaijmakers JM, Gardener BBM, Thomashow LS. Microbial populations responsible for specific soil suppressiveness to plant pathogens. Annu Rev Phytopathol 2002;40:309–48.

[60] Schlatter D, Kinkel L, Thomashow L, Weller D, Paulitz T. Disease suppressive soils: new insights from the soil microbiome. Phytopathology 2017;107:1284–97.

[61] Berg G, Rybakova D, Grube M, Köberl M. The plant microbiome explored: implications for experimental botany. J Exp Bot 2015;67:995–1002.

[62] Ellis JG. Can plant microbiome studies lead to effective biocontrol of plant diseases? Mol Plant Microbe Interact 2017;30:190–3.

[63] Berendsen RL, Pieterse CM, Bakker PA. The rhizosphere microbiome and plant health. Trends Plant Sci 2012;17:478–486.

[64] Suffert F, Sache I, Lannou C. Assessment of quantitative traits of aggressiveness in *Mycosphaerella graminicola* on adult wheat plants. Plant Pathol 2013;62:1330–41.

[65] Waalwijk C, Mendes O, Verstappen EC, de Waard MA, Kema GH. Isolation and characterization of the mating-type idiomorphs from the wheat septoria leaf blotch fungus *Mycosphaerella graminicola*. Fungal Genet Biol 2002;35:277–86.

[66] Buée M, Reich M, Murat C, Morin E, Nilsson RH, Uroz S, et al. 454 Pyrosequencing analyses of forest soils reveal an unexpectedly high fungal diversity: Research. New Phytol 2009;184:449–56.

[67] Caporaso JG, Lauber CL, Walters WA, Berg-Lyons D, Lozupone CA, Turnbaugh PJ, et al. Global patterns of 16S rRNA diversity at a depth of millions of sequences per sample. Proc Natl Acad Sci 2011;108:4516–22.

[68] Martin M. Cutadapt removes adapter sequences from high-throughput sequencing reads. EMBnet Journal 2011;17:10–2.

[69] Callahan BJ, McMurdie PJ, Rosen MJ, Han AW, Johnson AJA, Holmes SP. DADA2: High-resolution sample inference from Illumina amplicon data. Nat Methods 2016;13:581–3.

[70] Callahan B. DADA2 pipeline tutorial (1.8). https://benjjneb.github.io/dada2/tutorial.html (accessed March 20, 2019).

[71] Cole JR, Wang Q, Cardenas E, Fish J, Chai B, Farris RJ, et al. The Ribosomal Database Project: improved alignments and new tools for rRNA analysis. Nucleic Acids Res 2009;37:D141–5.

[72] Abarenkov K, Nilsson RH, Larsson K-H, Alexander IJ, Eberhardt U, Erland S, et al. The UNITE database for molecular identification of fungi – recent updates and future perspectives. New Phytol 2010;186:281–5.

[73] McMurdie PJ, Holmes S. phyloseq: An R package for reproducible interactive analysis and graphics of microbiome census data. PLoS ONE 2013;8:e61217.

[74] Paradis E, Schliep K. ape 5.0: an environment for modern phylogenetics and evolutionary analyses in R. Bioinformatics 2019; 35:526–8.

[75] Oksanen J, Blanchet FG, Friendly M, Kindt R, Legendre P, McGlinn D, et al. vegan: Community Ecology Package. R package version 2.5-4. https://CRAN.R-project.org/package=vegan (accessed March 20, 2019).

[76] Segata N, Izard J, Waldron L, Gevers D, Miropolsky L, Garrett WS, et al. Metagenomic biomarker discovery and explanation. Genome Biol 2011;12:R60.

[77] Kurtz ZD, Müller CL, Miraldi ER, Littman DR, Blaser MJ, Bonneau RA. Sparse and compositionally robust inference of microbial ecological networks. PLOS Comput Biol 2015;11:e1004226.

[78] Tipton L, Müller CL, Kurtz ZD, Huang L, Kleerup E, Morris A, et al. Fungi stabilize connectivity in the lung and skin microbial ecosystems. Microbiome 2018;6:12.

[79] Berry D, Widder S. Deciphering microbial interactions and detecting keystone species with co-occurrence networks. Front Microbiol 2014;5. doi:10.3389/fmicb.2014.00219.

[80] Csardi G, Nepusz T. The igraph software package for complex network research. https://pdfs.semanticscholar.org/1d27/44b83519657f5f2610698a8ddd177ced4f5c.pdf (accessed March 20, 2019).

[81] Shannon P, Markiel A, Ozier O, Baliga NS, Wang JT, Ramage D, et al. Cytoscape: a software environment for integrated models of biomolecular interaction networks. Genome Res 2003;13:2498–504.

